# mRNA initiation and termination are spatially coordinated

**DOI:** 10.1101/2024.01.05.574404

**Authors:** Ezequiel Calvo-Roitberg, Christine L. Carroll, Sergey V. Venev, GyeungYun Kim, Steven T. Mick, Job Dekker, Ana Fiszbein, Athma A. Pai

## Abstract

The expression of a precise mRNA transcriptome is crucial for establishing cell identity and function, with dozens of alternative isoforms produced for a single gene sequence. The regulation of mRNA isoform usage occurs by the coordination of co-transcriptional mRNA processing mechanisms across a gene. Decisions involved in mRNA initiation and termination underlie the largest extent of mRNA isoform diversity, but little is known about any relationships between decisions at both ends of mRNA molecules. Here, we systematically profile the joint usage of mRNA transcription start sites (TSSs) and polyadenylation sites (PASs) across tissues and species. Using both short and long read RNA-seq data, we observe that mRNAs preferentially using upstream TSSs also tend to use upstream PASs, and congruently, the usage of downstream sites is similarly paired. This observation suggests that mRNA 5’ end choice may directly influence mRNA 3’ ends. Our results suggest a novel “Positional Initiation-Termination Axis” (PITA), in which the usage of alternative terminal sites are coupled based on the order in which they appear in the genome. PITA isoforms are more likely to encode alternative protein domains and use conserved sites. PITA is strongly associated with the length of genomic features, such that PITA is enriched in longer genes with more area devoted to regions that regulate alternative 5’ or 3’ ends. Strikingly, we found that PITA genes are more likely than non-PITA genes to have multiple, overlapping chromatin structural domains related to pairing of ordinally coupled start and end sites. In turn, PITA coupling is also associated with fast RNA Polymerase II (RNAPII) trafficking across these long gene regions. Our findings indicate that a combination of spatial and kinetic mechanisms couple transcription initiation and mRNA 3’ end decisions based on ordinal position to define the expression mRNA isoforms.

## INTRODUCTION

The spatiotemporal control of gene expression is responsible for cell identity and function in all living species. In eukaryotes, RNA polymerase II (RNAPII) synthesizes pre-messenger RNAs (pre-mRNAs), which are processed by co-transcriptional mechanisms including transcription initiation and 5’ capping, splicing, and 3’ cleavage and polyadenylation (CPA) to create mature messenger RNA (mRNA). Each RNA processing step can be alternatively regulated, whereby different sites can be chosen during pre-mRNA synthesis to express a diverse set of mRNA isoforms from the same gene with variable transcript start sites (TSS), polyadenylation sites (PASs), or exonic compositions ^1^.

Mechanisms involved in RNA processing are regulated by disparate molecular machineries and subjected to different global regulatory constraints to drive tissue- or context-specific transcriptomes^2^. Thus, the regulation of exons across genes has historically been studied as distinct events that are regulated independently and contribute individually to the composition of full-length isoforms^3,4^. However, increasing evidence suggests cooperative regulation by RNA processing mechanisms across a gene^5–8^. Recent studies have found that factors involved in transcription, splicing, and CPA may have independent secondary roles in another step of RNA processing^9–13^. These investigations have mostly focused on connections between splicing and either transcription initiation or CPA, thus there is less known about direct connections between the regulation of transcription initiation and CPA.

Significant genetic space is devoted to regions involved in either transcription initiation or 3’ end processing. Greater than 70% of mammalian genes express alternative TSSs or PASs across cellular contexts^1,5,14^, with an average of 4-5 annotated alternative TSSs and/or PASs per gene^15,16^. Alternative TSSs and PASs influence the composition and length of coding sequences and 5’ and 3’ untranslated regions (UTRs), respectively, with impacts on RNA stability, localization, and translation efficiency^17–19^. The modulation of TSS and PAS usage is widely observed during proliferation, development, differentiation, and disease progression^20,21^. Overall, alternative mRNA terminal end usage (including both alternative start and end sites) underlies the majority of variation in isoform usage across cells and tissue types^5,22^. Understanding whether and how these decisions are coordinated across the gene will provide insight into the regulation of full-length isoforms, the proteins encoded by them, and effects on protein expression.

It is clear that there are many shared regulatory features between the 5’ and 3’ ends of genes. For instance, the same RNAPII phosphorylation events that are required for promoter-proximal transcription elongation are also required for productive elongation at the 3’ end of the gene^10,23,24^. Similarly, RNAPII elongation rates can influence both alternative TSS and PAS usage based on the position of sites within genes, such that there are global shifts towards either upstream or downstream TSSs^25–27^ or PASs^28–31^. Further, despite being separated by hundreds to thousands of nucleotides, decisions at the terminal ends of genes might also be spatially associated^32^. Recent high-resolution measurement of chromatin conformation found dynamic loops that are anchored at TSSs and extrude through the body of the gene^33,34^, suggesting that both two and three dimensional architecture may be important for mRNA synthesis. Together, these findings all hint at spatiotemporal connections between transcription initiation and CPA.

We set out to answer two fundamental questions in RNA biology: Are transcription initiation and 3’ end processing co-regulated and which mechanisms drive coordinated RNA processing? Here, through an analysis of genome-wide datasets across tissues, we find that genes tend to have similar numbers of alternative sites at both their 5’ and 3’ ends and there is a global bias towards coordinated usage of alternative TSSs and PASs based on their ordinal position within the gene. Our analyses suggest a “Positional Initiation-Termination Axis” (PITA) that governs the intrinsic choice of mRNA terminal ends independent of any tissue- or context-specific regulation. We describe the genome-wide relationships between mRNA terminal ends and characterize the mechanisms underlying their coupled regulation.

## RESULTS

The selection of mRNA terminal (5’ and 3’) ends is a highly regulated process that often involves a choice between multiple alternative sites and exons (**Fig. 1A**). To understand the extent to which isoform diversity in genes is driven by the usage of alternative terminal exons, we first systematically characterized the landscape of terminal exon usage. We previously developed the Hybrid-Internal-Terminal (HIT) index, a metric that uses RNA-seq data to identify and quantify the relative usage of alternative terminal exons across the genome^35^. We applied the HIT index framework to RNA-seq samples from eleven human tissues from the GTEx project^36^ and identified 29,622 and 35,985 alternative first and last exons, respectively, across all tissues. Surprisingly, we saw that the distributions of the number of first and last exons per gene were similar, where the number of genes that had one, two, three, or more first exons were similar to the number of genes that had only one, two, three, or more last exons, respectively (**Fig. 1B**). Genes with alternative first exons (AFEs) are significantly more likely than expected to also use alternative last exons (ALEs) (Hypergeometric Test p-value < 2.2 × 10^−16^; **Extended Data Fig. 1A**).

**Figure 1.**
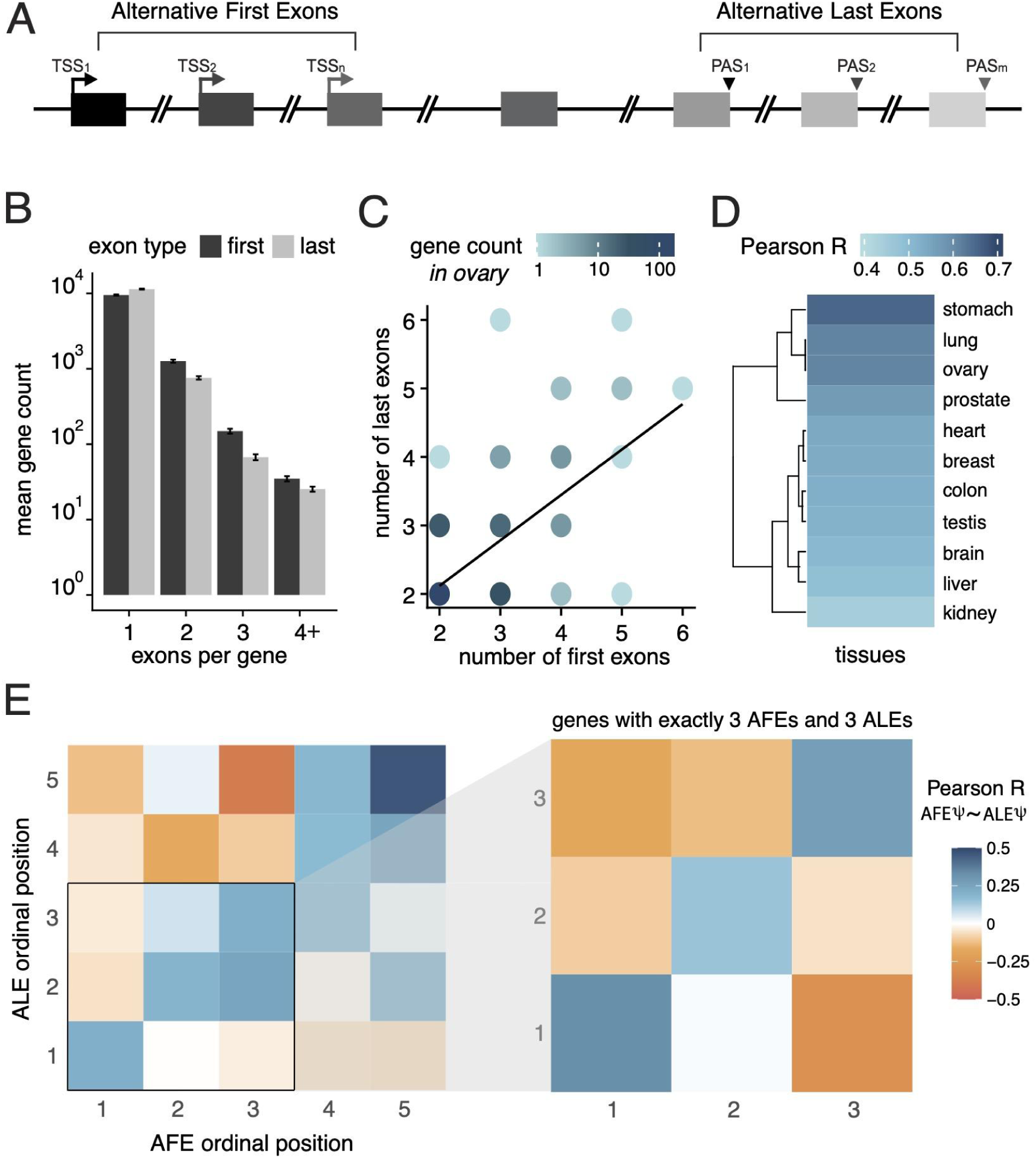
A positional relationship between sites of mRNA initiation and termination in human genes. **(A)** Schematic of a gene with alternative start and end sites. Alternative first exons (AFEs*, dark gray*) and their respective transcription start sites (TSSs) are shown on left, while alternative last exons (ALEs, *light gray*) and their respective polyadenylation sites (PASs) are shown on right. Exons are numbered by the order in which they appear in the direction of transcription (ordinal position). **(B)** The mean number of genes (*y-axis*) that express one, two, three, or at least four AFEs (*black*) or ALEs (*grey*). Error bars indicate standard error of the mean across tissues. **(C)** The correlation between the number of ALEs (*y-axis*) and AFEs (*x-axis*) used by each gene in a representative ovary tissue sample (n = 205, Pearson’s R = 0.76). Color gradient indicates the number of observations for each pairwise combination. (**D)** Correlations between the number of AFEs and ALEs across genes in 11 human tissues (color gradient indicates Pearson R)**. (E)** Heatmap of Pearson’s R values for pairwise correlations between the relative usage (Ψ) of AFEs and ALEs based on their ordinal position for genes with multiple first and last exons (up to 5; *left panel*, n = 5,559) and with exactly 3 AFEs and 3 ALEs (*right panel*, n = 227).

### Coordinated terminal exon usage based on ordinal position

This led us to speculate that genes might utilize similar numbers of first and last exons to jointly regulate the usage of terminal ends. Among genes with multiple AFEs and ALEs, there is a strong correlation between the number of first and last exons within a gene, such that genes using two AFEs are more likely to use two ALEs, genes using three AFEs often use three ALEs, *etc.* (Pearson R = 0.65 in ovary, p-value < 2.2 × 10^−16^ ; **Fig. 1C**). While the strength of this effect varies across tissues, all tissues analyzed showed strongly positive correlations (Pearson R > 0.4) between quantities of first and last exons identified despite tissue-specific regulation of alternative terminal end usage (**Fig. 1D**). The presence of similar numbers of alternative sites at both gene ends suggests the potential for regulatory coupling of transcription and CPA decisions based on gene organization, so we next asked whether the usage of first and last exons was associated with their position within the gene. As a baseline, genes that predominantly use a single first exon (measured by percent spliced in^37^ (PSI) > 0.95) also tend to use a single last exon (**Extended Data Fig. 1B**). More broadly, we found that the usage of terminal exons with similar ordinal positions was correlated across genes (**Fig. 1E**). Specifically, increased usage of the upstream most AFE (AFE 1) was associated with increased usage of the upstream most ALE (ALE 1), AFE 2 usage positively correlated with ALE 2 usage, *etc*. Concordantly, we observed that usage of AFE 1 negatively correlated with usage of ALE 2, indicating a complex coupling relationship based on the ordinal positions of the terminal exons within genes. Restricting the analysis to genes that use exactly three AFEs and three ALEs confirmed this ordinal coupling pattern (**Fig. 1E inset**), suggesting that not only are the number of terminal exons interconnected, but their usage is coupled as well. These findings indicate that gene architecture plays a pivotal role in determining coupling between terminal exons and thus regulating isoform diversity in genes.

### A positional initiation-termination axis that defines terminal exon coupling

The co-regulation of terminal exon usage could occur in one of two ways. Terminal sites may be directly coupled, whereby exons with similar ordinal positions are more likely to co-occur on the same mRNA molecules (as depicted in **Extended Data Fig. 1C**). However, it is also possible that they are indirectly coupled, such that exons with similar ordinal positions are independently favored across a population of mRNA molecules and thus appear to be correlated in short-read sequencing data (as depicted in **Extended Data** Fig 1D). To distinguish between these possibilities and identify specific genes with coupled terminal end usage, we analyzed long read ISO-seq data across human tissues and cells from the ENCODE Project^38,39^ and evaluated if terminal exons were coupled within the same molecules. Specifically, we wanted to quantify how often an mRNA that started at a particular transcription start site (TSS) was likely to terminate at a polyadenylation site (PAS) with a similar ordinal position. For example, after conditioning on full-length reads (Methods^40,41^; **Extended Data Fig. 2**), *MYO10* has 184 long mRNA reads in H9 cells, distributed across 3 TSSs and 2 primary PASs. 94% of the 128 reads that begin at the first TSS terminate at the first primary PAS, while 59% of the remaining 56 reads, that start downstream of the first TSS, terminate at the second primary PAS (**Fig. 2A**). Thus, long reads from *MYO10* support direct coupling of terminal exons based on their ordinal position, consistent with evidence from short-read RNA-seq data. These observations support the existence of a novel intra-molecular regulatory paradigm, which we call the “Positional Initiation-Termination Axis” (PITA), that underlies the coupling of transcription initiation and CPA decisions.

**Figure 2.**
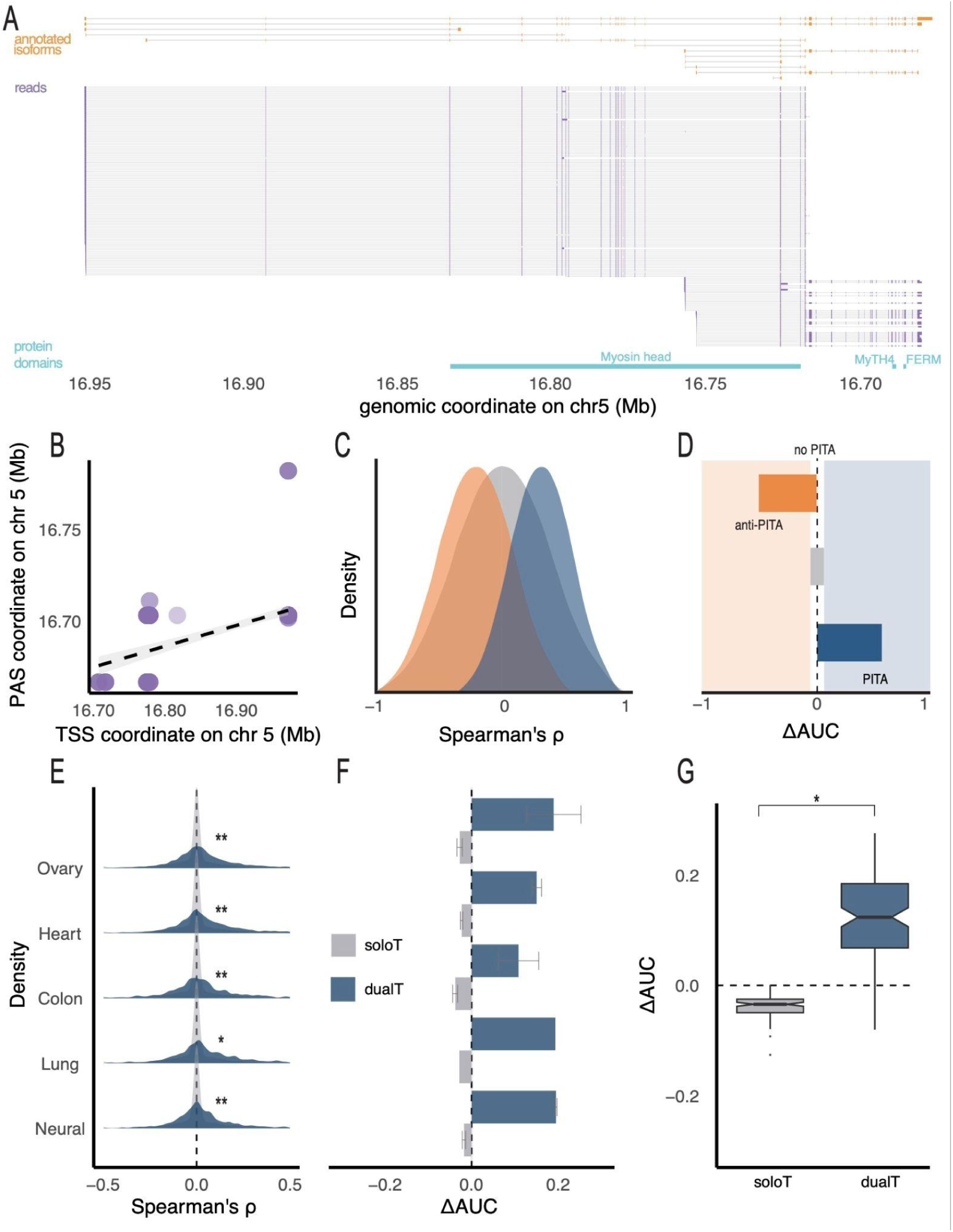
Positional coupling of TSS-PAS usage occurs within individual mRNAmolecules. (**A**) Annotated isoforms (*top*, *orange*), a subset of LRS reads (*middle*, *purple with introns in thin black lines*), and annotated protein domains (*bottom*, *light blue*) in H9 cells for *MYO10*. (**B**) Correlation between LRS read start (*x-axis*) and end coordinates (*y-axis*) for *MYO10* (Spearman’s ρ = 0.66). (**C**) Schematic of expected genome-wide distributions for Spearman’s ρ showing possible shifts towards negative correlations (*orange,* anti-PITA), positive correlations (*blue*, PITA) or an unbiased distribution (*gray*, no PITA) using fictional data (**D**) Schematic of expected ΔAUCs for the categories outlined in C. ΔAUC is defined as AUC_ρ>0_ - AUC_ρ<0_. Distribution of Spearman’s ρ **(E)** and the mean ΔAUC across samples **(F)** for solo termini genes (soloT, *gray*) and dual alternative termini genes(dualT, *blue*) in five human tissue types. K-S test * p-value < 10^−8^; ** p-value < 10^−16^ for **E**. (**G**) Distribution of ΔAUC values across 109 long-read sequencing samples for soloT (*gray*) and dualT genes (*blue*). T-test * p-value < 10^−16^.

To evaluate the extent to which intra-molecular coupling of ordinal terminal exons occurs globally, we calculated a Spearman ⍴ to look at the rank correlation between TSS and PAS positions across reads within a gene. *MYO10* has a Spearman ⍴ = 0.66 (**Fig. 2B**). Since rank corresponds to the ordinal position across terminal sites, positive correlations indicate ordinal coupling (PITA) while negative correlations indicate coupling of exons according to inverse ordinal position (anti-PITA) (schematic of possible distributions shown in **Fig. 2C**). We then used ΔAUC to quantify whether the distribution of Spearman ⍴s per sample shifted away being centered around 0 (Methods), where positive or negative ΔAUCs would again indicate genome-wide enrichments of PITA or anti-PITA genes, respectively (schematic of possible ΔAUCs shown in **Fig. 2D**).

When looking at genes that use multiple first exons and multiple PASs within a cell type (dual alternative termini (dualT)), we see significant enrichments of PITA genes across multiple human cells and tissues (shown for five tissues in **Fig. 2E-F**). We considered genes that use only one first exon or one PAS within each sample (solo termini (soloT)) as a control that accounts for variability in long read start and end coordinates and see that these genes consistently show weakly negative ΔAUCs, indicating our observations are not due to biases in the metric or identification of terminal ends from long reads. These trends are also not biased by read depth (**Extended Data Fig. 3A**), gene expression levels (**Extended Data Fig. 3B**), read length (**Extended Data Fig. 3C**), or other spurious correlations in the data (assessed with permutations, **Extended Data Fig. 3D**). Notably, the ΔAUCs are even more biased towards positive values when we calculated Pearson correlations using ordinal positions of sites rather than genomic coordinates of reads (**Extended Data Fig. 3E**-F) and the enrichment remains across different thresholds for calculating the ΔAUC (**Extended Data Fig. 3G**). On average, the distribution of PITA coupling across genes is more similar between replicates than non-replicates (**Extended Data Fig. 3H**). Finally, we see a similar enrichment of PITA genes when we use data from orthogonal experimental approaches (TIF-seq and cDNA-PCR datasets; **Extended Data Fig. 4**). Overall, we see that 80% of samples show evidence for greater than expected PITA coupling, with a 3-14% enrichment of PITA genes coupling across 109 ISO-seq samples from 47 tissues and cell types (**Fig. 2G**). Together, our results suggest that there is direct coupling of terminal exon usage and we can identify genes that are enriched for coupled terminal exon usage. This relationship is biased towards coupled usage of terminal exons with similar ordinal positions in a gene (**Fig. 3A**).

**Figure 3.**
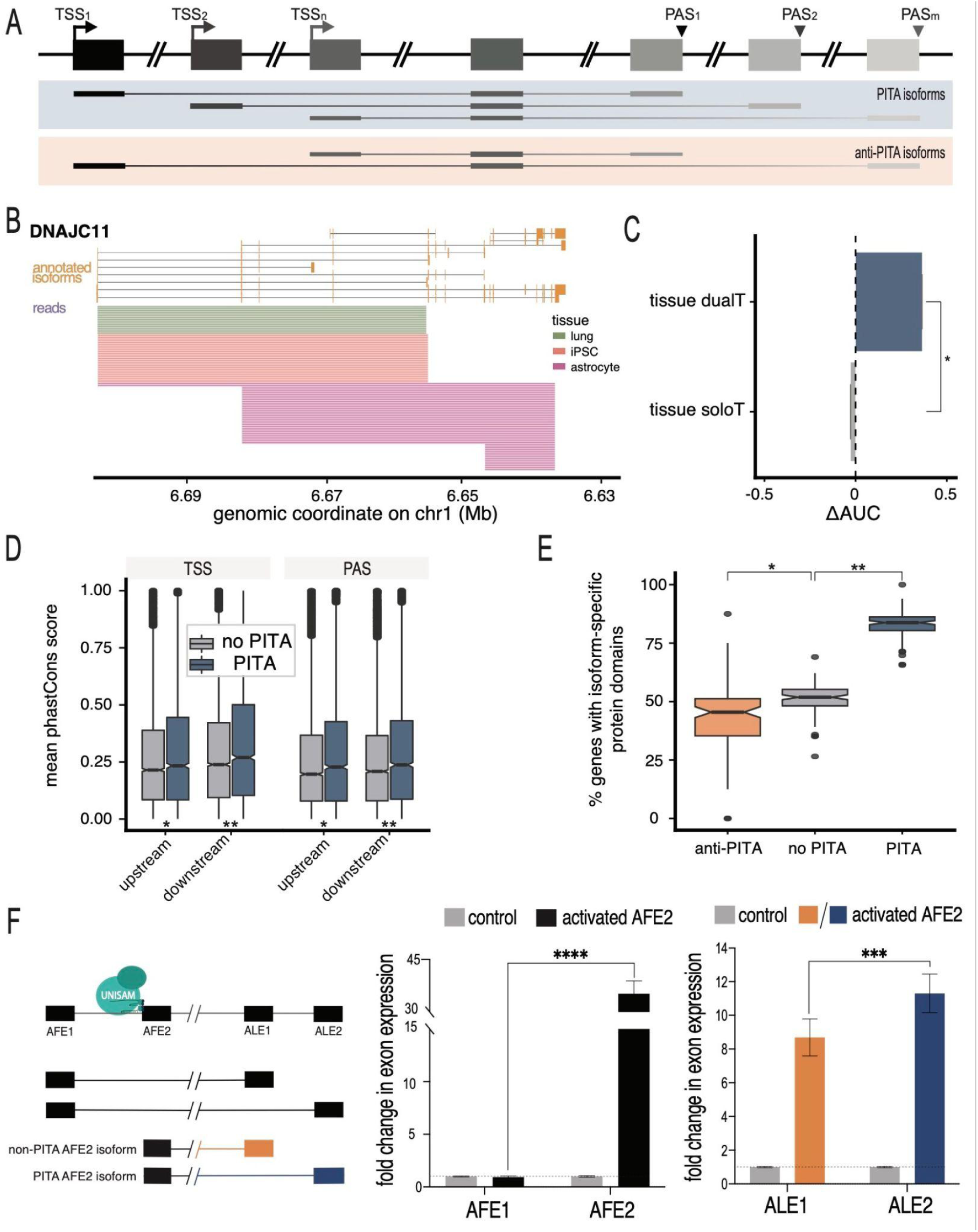
Transcription start sites directly regulate transcript end site usage to govern the expression of functionally distinct isoforms. (**A**) Schematic of mRNA isoforms based on PITA classification. PITA isoforms preferentially use TSSs and PASs that are ordinally similar (l*ight blue*), while anti-PITA isoforms preferentially use ordinally different TSSs and PASs (*light orange*). (**B**) Annotated mRNA isoforms (*orange*) and a randomly subsampled proportion of reads for *DNAJC11* in lung (green), iPSCs (pink), and astrocytes (*purple*). (**C**) ΔAUC values for genes with dual alternative termini across tissues. Error bars represent standard error across 100 samples of reads across tissues. T-test * p-value < 10^−16^. (**D**) Conservation scores (mean phastCons score, *y-axis*) in a 400nt region around each terminal sites of the two most highly expressed isoforms for genes with dual alternative termini. K-S test * p-value < 10^−3^; ** p-value < 10^−7^. (**E**) Percentage of dual alternative termini genes (*y-axis*) whose isoforms overlap different annotated protein domains. T-test * p-value < 10^−7^; ** p-value < 10^−16^. (**F**) CRISPR activation of the second AFE of *LEPR* in HEK293 cells. Schematic (*left*) of LEPR possible isoforms and UNISAM-gRNA activation complex target site upstream of *LEPR* AFE2. Fold change in AFE expression measured by qRT-PCR *(middle panel)* upon treatment of cells with UNISAM-gRNA targeting AFE2 *(black)* or an empty vector control *(gray).* Fold change in expression of ALE1 (orange, non-PITA) or ALE2 (blue, PITA) upon activation of AFE2, measured by qRT-PCR *(right panel).* *** p-value < 10^−3^ **** p-value < 10^−4^.

### PITA is independent of tissue-specific mRNA regulation

mRNA terminal ends are often regulated across tissues, with abundant tissue-specific usage of both TSSs and PASs. Indeed, 83% of genes expressed in the long-read dataset are regulated in a tissue-specific manner, such that they use distinct isoforms across tissues. It is also known that tissue- or context-specific regulation of alternative TSSs or PASs results in global shifts towards preferential usage of upstream or downstream sites^17,42–46^. And even among dualT genes there is usually one predominant isoform expressed, likely due to tissue-specific regulation of isoform usage and gene expression. When we control for this differential isoform usage and gene expression, the enrichment of PITA genes increases (9-23% enrichment; **Extended Data Fig. 3B**). Thus, PITA coupling within dualT genes is likely not a by-product of global tissue-specific signatures, but an independent driver of full-length isoform usage. To investigate this regulatory paradigm further, we asked whether genes that show strong tissue-specific regulation of terminal sites also show PITA coupling across tissues. Specifically, we looked at genes that use unique termini within tissues, but use alternative TSSs and alternative PASs across tissues (tissue-dualT). For example, *DNJC11* expresses distinct TSSs and PASs across lung, iPSC, and astrocyte cells (**Fig. 3B**). However, across cell types there is a significant correlation of *DNJC11* TSS and PAS usage, with the expressed isoforms showing a preference for PITA coupling (Spearman ⍴ = 0.92). Using a tissue-ΔAUC metric, where Spearman ⍴s are calculated using reads across tissues for tissue-dualT genes (Methods), we see an enrichment of PITA coupling that is stronger than that observed in most individual tissues (17-19% enrichment; **Fig. 3C**). This observation suggests that PITA coupling is a pervasive phenomenon where the coordination of transcript starts and ends might be influenced by gene architecture even when isoforms are tissue-specifically regulated or constitutively expressed within a cell type.

The preferential usage of PITA isoforms may enable the regulation of RNA or protein features related to the positioning of elements within a gene. For instance, if PITA coupling allows for the regulation of isoforms with specific functional significance, PITA genes may show conserved features specifically at the terminal ends of transcripts. Consistent with this, the upstream and downstream TSS and PAS regions for dualT genes with PITA coupling are more conserved than similar regions for non-PITA or anti-PITA dualT genes (**Fig. 3D**). Furthermore, genes expressing PITA isoforms more often encode for a greater variety of protein domains than anti-PITA or non-PITA genes (Methods, **Fig. 3E**), despite all of these genes containing similar numbers of domains on average (**Extended Data Fig. 5A**-B). For instance, *MYO10* PITA isoforms starting at the upstream TSS are able to encode a myosin head domain, while PITA isoforms starting at the downstream PAS are able to encode MyTH4 and FER domains (**Fig. 2A**, *bottom track*). Together, these observations suggest that PITA coupling may help to shape mRNA function or proteome diversity through coordinated exon usage across the terminal ends of an isoform. Notably, we do not see either increased conservation or protein domain diversity for genes with inter-tissue PITA coupling (**Extended Data Fig. 5B**-C), suggesting that all tissue-dualT genes may already be under stronger functional constraints due to cell-type specific isoform regulation^18,20,47^.

### Alternative first exon usage directly influences last exon usage

These correlative analyses suggest the possibility of a causal relationship between the ordinal coupled usage of alternative TSSs and alternative PASs. To test causation, we used a dCas9-CRISPR activation system to specifically activate the second AFE of *LEPR* in HEK293 cells^48^ (Methods; **Fig. 3F; Extended Data** Fig 5D-E). We first confirmed that the expression of AFE2 increases 35-fold upon dCas9-mediated activation (T-test p-value < 1 × 10^−6^), while the relative expression of AFE1 does not significantly change (T-test p-value = 0.11). AFE2 activation led to increased relative expression of both ALE1 and ALE2, however ALE2 was significantly more activated than ALE1 (1.3-fold increase in relative activation, T-test p-value = 0.0002). These results confirm that AFE usage influences ALE regulation and that there is a direct relationship between the usage of terminal exons in an ordinal fashion.

### Gene length drives PITA coupling

PITA coupling suggests that the architectures of genes are configured to enable co-regulation of transcript start and end sites. This coupling involves long distance coordination across transcription initiation and polyA sites that are often separated by thousands to hundreds of thousands of base pairs in DNA space and hundreds to thousands of nucleotides in pre-mRNA space. Thus, we first evaluated whether the DNA or RNA distance between these sites was correlated with the strength of PITA coupling across genes. PITA coupling is more likely to occur in longer genes, defined as the total genomic length between the upstream most expressed TSS and downstream most expressed PAS (**Fig. 4A**). However, PITA coupling is not associated with the pre-mRNA length of the resulting isoforms (**Fig. 4A**) or the mRNA lengths of the isoforms expressed (average read length, Methods; **Extended Data Fig. 6A**). This observation suggests that PITA coupling may be driven by transcriptional or co-transcriptional mechanisms and regulatory decisions.

**Figure 4.**
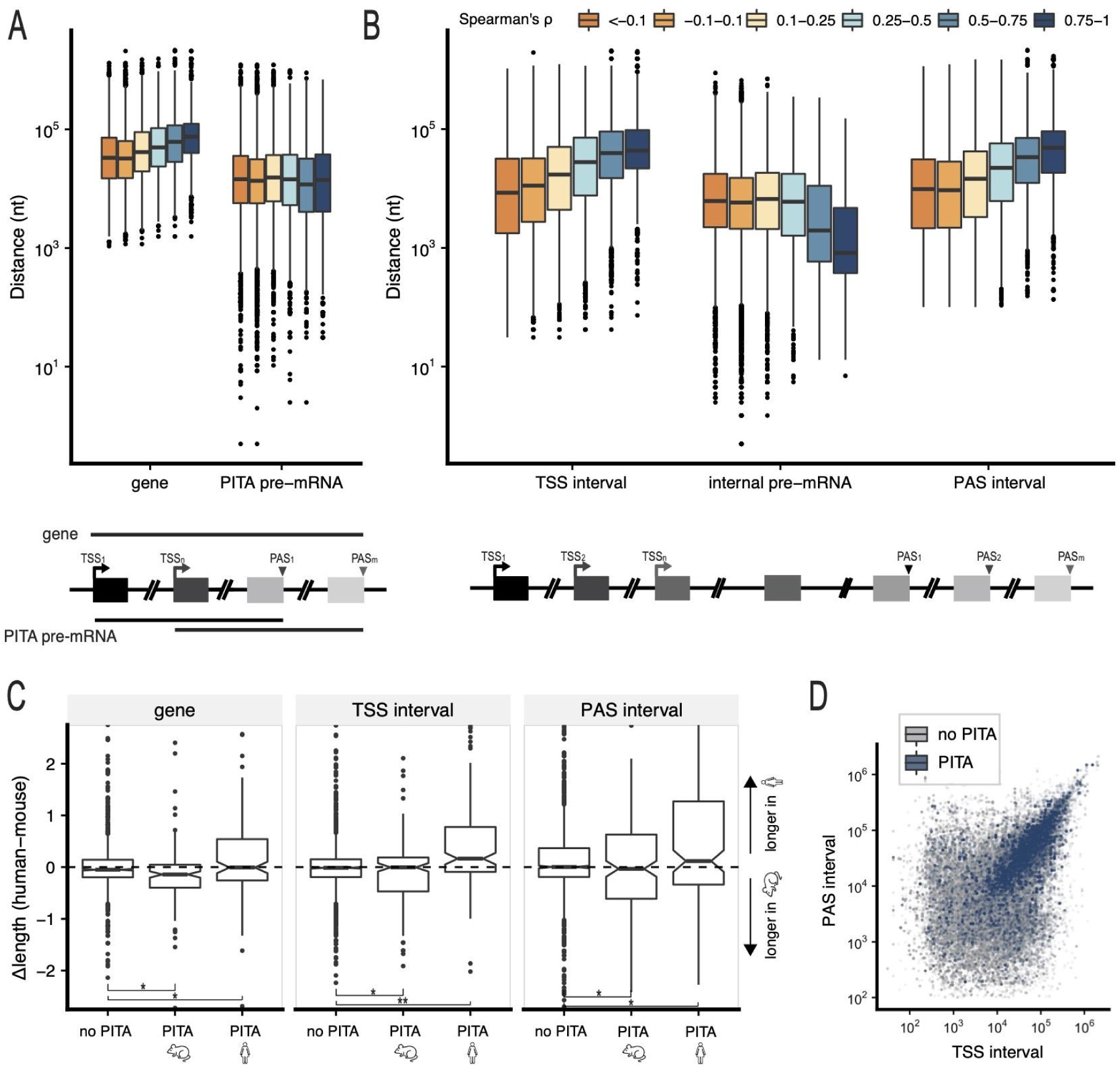
PITA occurs more often in longer genes. (**A**) Distribution of the lengths (*y-axis*) of dual alternative termini genes (*left*) and PITA pre-mRNAs (*right*) for genes within bins of Spearman’s ρ values (*colors*). (**B**) Distribution of the maximum distances (*y-axis*) between TSSs (*left*), the downstream-most TSS and upstream-most PAS (internal pre-mRNA, *middle*), and PASs (*right*) for dual alternative termini genes within varying bins of Spearman’s ρ values (*colors*). (**C**) Distribution of the difference in feature length for human and mouse orthologs (*y-axis*) of genes that are not PITA in either species, PITA only in mice or PITA only in humans. To account for global differences in gene lengths between species, distances were first normalized by the mean distance within each species for each feature. K-S test * p-value < 10^−3^; ** p-value < 10^−7^. (**D**) Correlation between the TSS intervals (*x-axis*) and PAS intervals (*y-axis*) for dual alternative termini genes non-PITA genes (Spearman ρ < 0.3, *gray*, Pearson R = 0.59) and PITA genes (Spearman ρ >= 0.3, *blue*, Pearson R = 0.82).

A gene can be broken down into many regions, all of which contribute independently to the total length of a gene. To understand how the length of a gene may influence PITA coupling, we calculated lengths of the regions involved in transcriptional regulation (interval between TSSs), only involved in splicing regulation (interval between downstream most TSS and upstream most PAS), and involved in cleavage and polyadenylation regulation (interval between PASs). We find that both the TSS and PAS intervals are strongly associated with PITA, with longer intervals between alternative TSSs and alternative PASs leading to stronger PITA coupling (**Fig. 4C**). However, the length of the internal pre-mRNA interval is not associated with the strength of PITA coupling. To understand whether the TSS and PAS intervals were directly associated with PITA regulation, we used a comparative genomics approach and treated evolutionary changes as natural perturbations that vary gene length. Using ISO-seq data from mouse tissues matched to the human tissues analyzed (ENCODE^38,39^, Methods), we identified species-specific PITA genes as described above (**Extended Data Fig. 6B**-C). We find that genes only showing PITA coupling in human cells are, on average, significantly longer in the human genome than in the mouse genome and, correspondingly, mouse-specific PITA genes are significantly longer in the mouse genome (Kolmogorov-Smirnov test p-value < 0.001; **Fig. 4E, Extended Data Fig. 6D**). Reassuringly, genes that show no PITA coupling in either species do not vary in gene lengths. This pattern of species-specific PITA genes having longer genomic intervals in the PITA species also holds for TSS and PAS intervals. The opposite is also true, whereby genes with the largest difference in gene length or TSS and PAS interval lengths between species are enriched for species-specific PITA genes in the species with longer lengths (**Extended Data Fig. 6E**). Together, these observations suggest that the strength of PITA coupling is directly associated with longer genomic distances.

Since PITA coupling shows a relationship with both TSS and PAS intervals, we wanted to understand whether certain genes were more likely to show an association with TSS intervals and others with PAS intervals. Given our initial observation that genes are more likely to have similar numbers of first and last exons and a correlation between gene length and terminal interval length (**Extended Data Fig. 7 A-B**), we first asked whether genes are also more likely to have similar transcription and CPA region intervals. While TSS and PAS intervals are weakly correlated globally (Pearson R = 0.59, p-value < 2.2 × 10^−16^), this correlation is even stronger for PITA genes (Pearson R = 0.82, p-value < 2.2 × 10^−16^; **Fig. 4D**). Strikingly, the change in TSS interval length between human and mouse is also significantly correlated to the change in inter-species PAS interval length for human- and mouse-specific PITA genes (**Extended Data Fig. 7C**). Thus, PITA genes are jointly influenced by TSS and PAS interval ranges. Together, these observations suggest that there are selective pressures to coordinate the length of genomic regions that are involved in the regulation of alternative transcription and CPA.

### Distinct promoter regions and 3D architecture govern PITA coupling

The association between PITA coupling and the lengths of genomic intervals suggests that PITA may be driven by a mechanism constrained by the two- or three-dimensional organization of genes, including the relative positioning of regulatory elements at both terminal ends of a gene^34^. This paradigm is less likely to involve independent or coordinated recruitment of particular regulatory factors, since such models would not be expected to show gene length-dependence. Instead, we hypothesized that the paired regulatory intervals at gene terminal ends might govern structural or 3D-conformational regulation of mRNA isoform production. We used ultradeep micro-C data^49,50^ to investigate whether there is any evidence for specific localized physical interactions between the terminal ends of PITA or non-PITA genes, and/or whether PITA genes display more global differences in chromatin conformation at the level of chromatin domains and domain boundaries, as compared to non-PITA genes (Methods).

Previously, it has been shown by Hi-C that active TSSs and PASs display insulation^32^: these loci prevent long-range chromatin interactions across them, leading to the formation of structural chromatin domains containing active genes from start to end, and demarcated by insulating boundaries^32,34,51,52^. Consistently, here we also found that both non-PITA and PITA genes form structural domains defined across the entire gene, with insulation boundaries at TSS and PAS, in both human embryonic stem (H1-ES) and foreskin fibroblast (HFF) cells (**Fig. 5A, Extended Data Fig. 8A**-B**, Extended Data Fig. 9A**). Insulation at PASs appears to be less precisely positioned – likely because PASs are less well defined and more variable than TSSs between cells^29,53^ – and therefore more difficult to capture at the same resolution as TSSs, as shown before^52^. While non-PITA genes predominantly express the longest pre-mRNA isoforms, and at the chromatin level formed a single structural domain, PITA genes express more diverse isoforms using both upstream and downstream TSSs and PASs (**Extended Data Fig. 10A**-B). In line with this, PITA genes often have evidence for formation of multiple overlapping structural domains whose boundaries are defined by relationships between the upstream TSS - upstream PAS and downstream TSS - downstream PAS pair (**Fig. 5A, Extended Data Fig. 8A**), suggesting the presence of chromatin substructures within PITA genes that may govern the relationship between mRNA initiation and termination and isoform expression.

**Figure 5:**
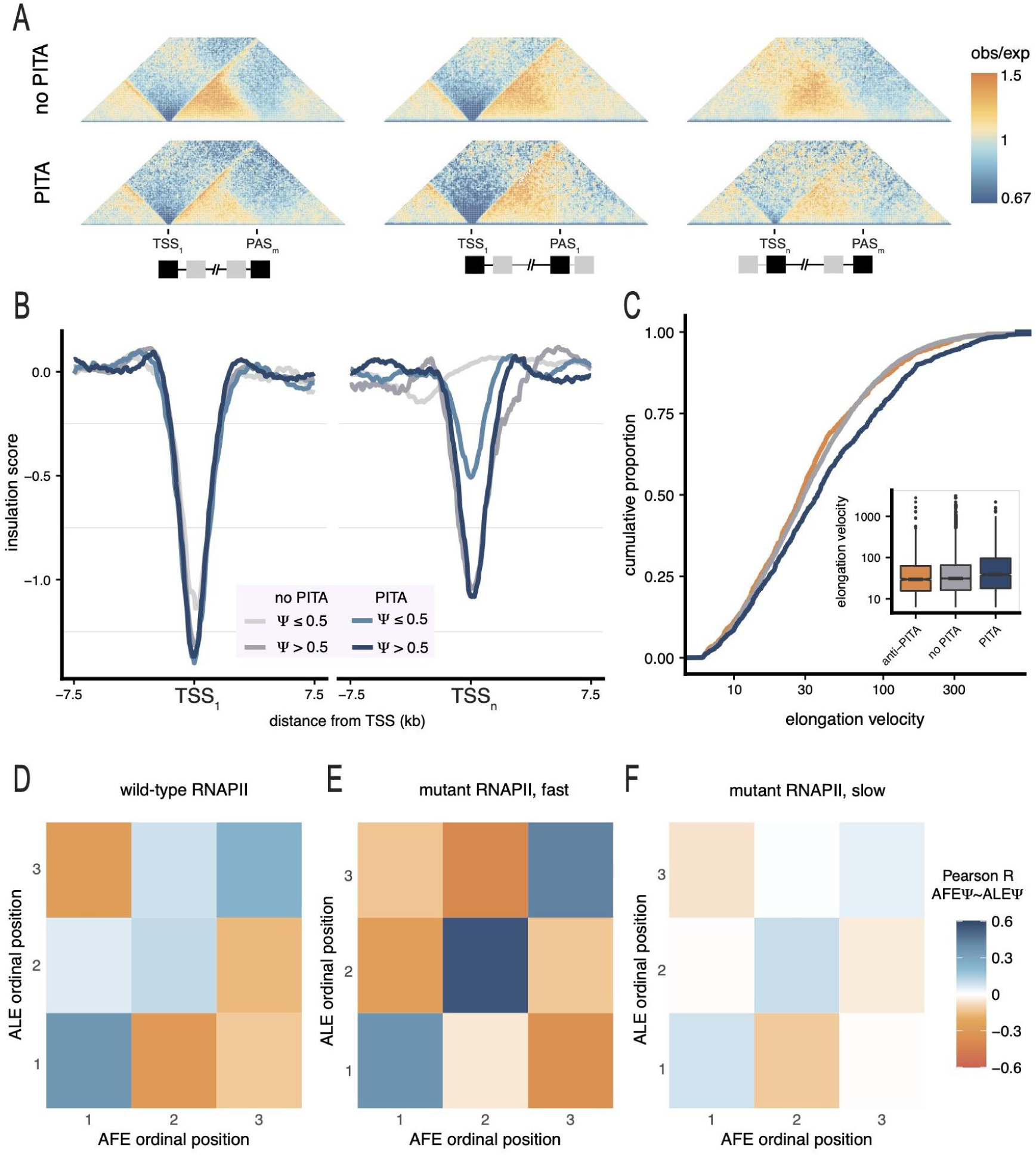
RNA polymerase II elongation rate is associated with PITA. **(A)** Aggregated and scaled micro-C maps for TSS1-PASn, TSS1-PAS1, and TSSn-PASm pairs at 500 bp resolution in H1-ESCs, where the color scale indicates low (depletion of interactions, *blue*) to high (enrichment of interactions, *orange*) microC signal enrichment, where each figure pixel reflects a paired interaction. **(B)** Average insulation profiles in H1-ESCs at 250 bp resolution in the +/-7.5kb window around upstream TSS_1_ (*left*) and downstream TSS_n_ (*right*) for non-PITA (*grey scale*) and PITA (*blue scale*) genes. Shades indicate relative TSS usage (measured by PSI values). (**C**) Cumulative distribution of elongation velocities in K562 cells for anti-PITA (*orange*), non PITA (*grey*) and PITA genes (*blue*), considering only dual alternative termini genes. Inset shows boxplots of the full distributions. K-S test * p-value < 0.05; ** p-value < 10^−15^. Heatmaps of Pearson’s R values for the pairwise correlations between the relative usage (Ψ) of a gene’s AFEs and ALEs based on their genomic order in HEK293 cells expressing **(E)** WT RNAPII, (n = 49), **(F)** fast elongating RNAPII (*R749H* mutation, n= 69) and **(G)** slowly elongating RNAPII (*E1126C* mutation, n= 62). All heatmaps show pairwise correlations for genes expressing exactly 3 AFEs and 3 AFEs.

While upstream TSSs consistently display insulation regardless of PITA categorization, downstream TSSs more clearly show aggregate evidence for insulation in PITA genes. This observation led us to hypothesize that alternative TSSs may have distinct regulatory patterns in PITA genes. For upstream TSSs in both PITA and non-PITA genes, insulation is accompanied by promoter-defining marks such as chromatin accessibility, H3K4me3, and CTCF binding (**Extended Data** Figs. 8B and 9A), regardless of the absolute or relative expression of this TSS (**Fig. 5B**). However, the presence of insulation and promoter-defining marks at downstream TSSs is correlated with the relative usage of the TSS. Specifically, major TSSs, including downstream TSSs (PSI > 0.5) have equally strong insulation in both non-PITA and PITA genes, while minor TSSs have insulation in PITA genes but not in non-PITA genes (**Fig. 5B and Extended Data** Figs. 8C). Finally, PITA genes also appear to have an additional insulation sub-domain across the TSS interval, bounded by the upstream most and downstream most TSS (**Extended Data** Figs. 8D and 9B). Together, these observations suggest that alternative TSSs in PITA genes may be regulated by distinct promoters (rather than a shared promoter region) and their alternative and coordinated terminal usages are reflected in distinct 3D genetic architectures.

### RNAPII elongation rates mediate the relationship between mRNA ends

The presence of distinct promoter regions across alternative TSSs in PITA genes prompts the exploration of additional molecular mechanisms governing productive transcription through this complex regulatory landscape. It is possible that the overlapping chromatin substructures within PITA genes may influence or be influenced by the recruitment or trafficking of RNAPII across the gene body. In particular, since PITA genes are significantly longer than non-PITA genes, the rate at which they are transcribed might be specifically regulated to ensure proper RNAPII processivity and recycling. We used the ratio of TT-seq (quantifying transcribed RNA) to mNET-seq (quantifying actively transcribing RNAPIIs) data in K562 cells to calculate RNAPII elongation velocities, as described before^54,55^. Consistent with our hypothesis, we see that PITA genes have significantly faster RNAPII elongation velocities relative to both non-PITA and anti-PITA genes (Methods, Kolmogorov-Smirnov test p value < 1 × 10^−15^, **Fig. 5C**). However, elongation rates are faster for longer genes, as has been observed previously^56^, and PITA genes are not transcribed faster on average than than expected based on their length (**Extended Data Fig. 10C**). Regardless, the relationship between elongation speed and gene length suggests a relationship between mRNA isoform expression and gene length such that, for each gene, RNAPII can only productively elongate for a given distance.

For long PITA genes, RNAPII molecules starting in upstream or downstream TSSs may efficiently transcribe far enough to use upstream or downstream PASs, respectively, purely by virtue of their faster elongation rates. To test this, we analyzed publicly available RNA-seq data from human cells overexpressing RNAPII mutants that elongate either slower or faster than wild-type RNAPII, paired with a control line (**Fig. 5D-F**)^57^. While the fast elongating RNAPII strengthened the enrichment of PITA coupling (**Fig. 5E**), the slow RNAPII markedly disrupted the correlated usage of ordinally similar terminal exons (**Fig. 5F**). This observation was consistent in an edited mouse cell line expressing the slow RNAPII mutation (**Extended Data Fig. 10D**)^58^. This suggests that RNAPII elongation rates are crucial for coupling TSS-PAS interactions in long PITA genes, ensuring sustained RNAPII processivity to reach longer range PASs. Together, our observations indicate that a combination of spatial and kinetic mechanisms contribute to a relationship between pre-mRNA ends and ultimate expression of mRNA isoforms (**Fig. 6**).

**Figure 6:**
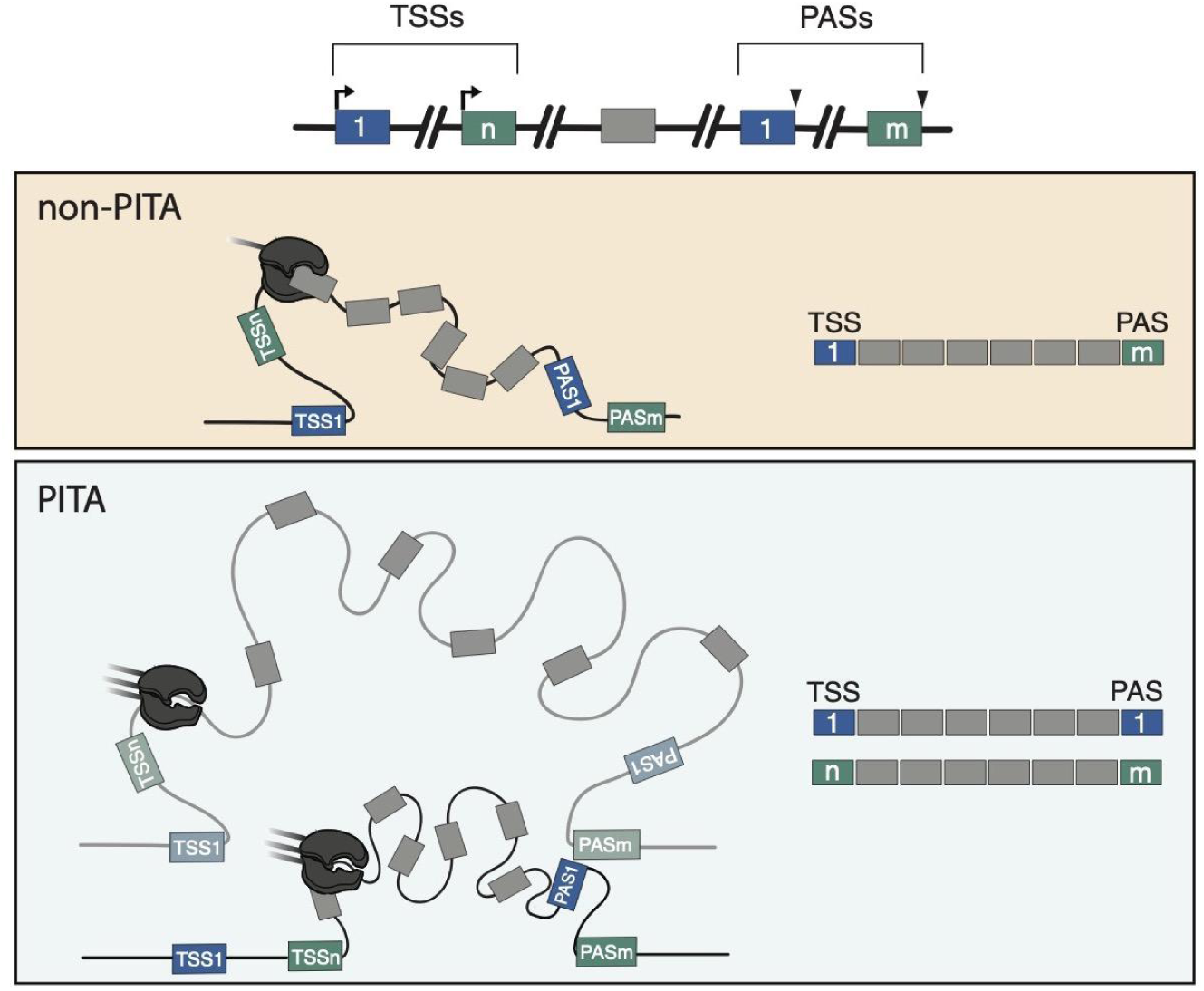
Spatial and kinetic mechanisms drive the ordinal coupling of mRNA initiation and termination in PITA genes. Schematic representations of representative genes. Transcription start (TSS) and polyadenylation (PAS) sites are identified by ordinal position 1 (*blue*) through ordinal position n/m (*green*). Internal exons are gray. Top panel depicts a non-PITA gene, which tends to be shorter, have average RNAPII elongation rates, and classic gene insulation across the gene body. Bottom panel depicts a PITA gene,which tends to be longer, have faster RNAPII elongation, and chromatin sub-domains bounded by ordinally coupled TSS and PAS pairs. Dominant mRNA isoforms are shown on the right.

## DISCUSSION

Here, we observe the coordinated usage of mRNA initiation and termination sites within a gene. This is consistent with a recent report in *Drosophila* tissues and human organoids showing that a dominant promoter region can be preferentially associated with usage of one or multiple polyadenylation sites^8^. However we find little overlap between PITA genes and genes with dominant promoters (**Extended Figure 4**), suggesting the potential for two non-mutually exclusive regulatory paradigms influencing the co-regulation of mRNA initiation and termination. We find that PITA coupling between TSSs and PASs is correlated through combinations of gene length and chromatin conformation, and is mechanistically driven by RNAPII elongation rates (**Fig. 6**). These molecular features act in concert to create a positional dependence in coordinated exon usage such that the isoforms starting at the upstream TSS favor usage of an upstream PAS and isoforms starting at a downstream TSS favor usage of a downstream PAS, defining a intrinsic positional initiation-termination axis (PITA) governing intrinsic isoform patterns in lieu of specific regulation.

Widespread coupling among different RNA processing events has been previously documented. Preliminary evidence showed intriguing associations between TSS, internal exons, and PAS using both short- and long-read datasets^1,2,5,59^. Biochemical and genetic evidence indicate that components of the splicing and CPA machinery, as well as auxiliary regulatory factors, are loaded onto the C-terminal domain (CTD) of RNAPII at the time of transcription initiation to facilitate co-transcriptional RNA-processing^12,60,61^. These factors may even moonlight as regulators of secondary processes, with evidence that transcription factors can regulate splicing^62,63^, splicing factors can locally recruit transcription factors^64,65^, and spliceosome components can influence the usage of premature PASs^9^. In line with these biochemical observations, studies have found connections between skipped exons and either TSSs or PASs over evolutionary timescales^6,66^, cancer and cellular transitions^31^, and across tissues^1,8^. Our observations of direct TSS-PAS coupling adds to this complex and coordinated regulatory landscape driving isoform usage. While preliminary evidence points to the possibility of interconnections between skipped exon usage and TSS-PAS co-regulation^8^, it remains to be seen to what extent PITA coupling also drives the composition of internal exons in an isoform.

Despite all this work, it has remained unclear what molecular forces drive these correlations. We show genomic evidence for gene architecture, chromatin conformation, and kinetic mechanisms all playing a role in PITA coupling. For non-PITA genes, we see chromatin architecture that defines an insulation domain encompassing the entire gene, consistent with previous observations and literature that suggests the involvement of chromatin domains in constraining RNAPII directionality^67^. Concurrently, there is preferential expression of the longest pre-mRNA isoform in non-PITA genes. However, in PITA genes, the picture is more complex, with multiple sub-domains bounded by ordinally coupled TSS-PAS pairs and inter-promoter interactions. PITA genes are also among the longest genes, which tend to have faster RNAPII elongation rates. Given that a slower RNAPII disrupts PITA coupling, our results align with the “window of opportunity” model for RNA processing, which indicates the existence of an optimal RNAPII elongation rate for RNA processing events^57^. This optimal rate appears to be crucial in enabling RNAPII to productively elongate until it reaches the coupled PAS and may especially play a role in longer genes where RNA processing sites are separated by longer genomic distances. However, under this model, it is surprising that isoforms initiating at a downstream TSS do not equally use either upstream or downstream PASs as they emerge from the RNAPII exit channel. However, evidence of an insulation sub-domain bounded by the downstream TSS and downstream PAS that appears to only exist in PITA genes suggests that spatial interactions may promote the preferential coupling of these sites.

Our results indicate that spatial and kinetic mechanisms jointly influence the coupled regulation of mRNA processing events across a gene. However, our analyses are unable to infer any causality or directionality between these processes - it is possible that chromatin conformations influence RNAPII processivity, or visa versa. Perhaps more likely, there is a dynamic interplay between both to maintain an efficient transcriptional environment and PITA coupling is a by-product of these regulatory fluctuations. Several pieces of evidence point to this possibility. Insulated sub-domains may structure chromatin to place specific regions within cellular sub-compartments enriched for transcription and/or splicing machinery, which recent work has indicated may promote RNAPII processivity^33,68–71^. This 3D architecture may be especially important for proper CPA, since it would bring RNAPII at the end of the gene in contact with kinases that phosphorylate RNAPII to initiate transcription and elongation. The same phosphorylation marks are also necessary for CPA, but it has been seen that phosphorylation levels diminish across a gene^10^, potentially setting an intrinsic limit on the productive elongation distance of RNAPII. An interplay between 3D and kinetic control of transcription elongation, as seen for PITA coupling, may be especially important for the transcription of longer genes and expression of mature isoforms. Finally, there is evidence for direct involvement of transcriptional enhancers and elongation factors contributing to the regulation of alternative PASs by binding to upstream enhancer elements^72^ or playing a role in recruiting PAS machinery^12^. Together, our description of coupled transcription and CPA choices sheds light on the mRNA consequences of these mechanistic features.

Genes with PITA coupling have, on average, significantly longer genomic lengths. Notably, a change in gene length across evolutionary time is associated with species-specific PITA coupling. Mouse genes showing mouse-specific PITA coupling are significantly longer than their human orthologs. It is unclear, however, whether PITA coupling arises due to a change in genomic distances or vice versa. Interestingly, we found a strong association between a species-specific increase in both TSS and PAS interval lengths and the likelihood of species-specific PITA coupling, suggesting that selective pressures may constraint the lengths of gene regions involved in regulating alternative transcription initiation and 3’ end site usage. Recent evidence indicates that longer and more highly expressed genes in human cells are more likely to form cohesin-mediated extrusion loops that are bounded by transcription start and end sites^33^. Thus, it is possible that there is a minimum genomic interval required between TSSs and between PASs to establish paired insulation across TSSs and PASs in mammalian genomes. However, it is still unknown whether the first evolutionary determinant of PITA coupling is the regulation of terminal site usage, the distance between terminal sites, or the dynamics of RNAPII in the region.

Finally, we observe a stronger enrichment of PITA coupling across tissues than across isoforms within the same sample. This is consistent with the established importance of tissue-specific promoter and cleavage site usage and may aid in the diversification of isoform expression. We see that PITA coupling is enriched in genes for which usage of ordinally coupled mRNA terminal ends enables the expression of a more diverse set of protein domains. This raises the possibility that PITA coupling might be an intrinsic regulatory paradigm created by the 2- and 3-D architecture of genes and RNAPII elongation speeds across long genes, while non-PITA coupling might be actively regulated through mechanisms like paired insulation across gene starts and ends to allow for tissue-specific isoform expression. Alternatively, PITA coupling might naturally occur in all genes, but the formation of insulation domains across an entire gene in non-PITA genes disrupts this process.

PITA coupling represents paired mRNA 5’ and 3’ end usage that influences ultimate isoform compositions, adding complexity to the promoter-centric view of mRNA isoform regulation. We found that elements of gene architecture and RNA polymerase dynamics play key roles in shaping mRNA diversity. This intricate interplay of gene regulatory mechanisms may be further modulated by tissue-specific factors that increase the specific usage of terminal sites. Understanding these and additional regulators of coupled mRNA terminal site usage will offer a deeper understanding of how cellular context fine tunes isoform expression patterns and contributes to cellular identity.

## METHODS

### Short-Read RNA sequencing Analysis

RNA-sequencing data across 11 human tissues from different individuals (2 replicates per tissue) from GTEx Version 8 were downloaded through dbGaP under study accession number phs000424.v8.p2. RNA-seq data from RNAPII mutant cell lines was downloaded from the NCBI Gene Expression Omnibus (GSE63375). RNA-seq data from mouse data was downloaded from the NCBI Gene Expression Omnibus (GSE127741). A detailed description of short read sequencing samples used can be found in Supplementary Table 1. For all short-read sequencing data, reads were mapped to genome assembly GRCh38.p14 for human or GRCm38.95 for mouse, using STAR^73^. Resulting bam files were processed using the HIT index pipeline to classify and quantify first and last exon usage^35^. For exon classification, we utilized conservative parameters on all short read data analyzed, with a HIT_identity_parameters.txt file as follows: *# |HITindex| threshold for calling terminal exons [0.0, 1.0] HITterminal 1.0, # |HITindex| threshold for calling hybrid exons [0.0, 1.0] HIThybrid 0.5, # bootstrapping p-value threshold for HITindex significance [0.0, 1.0], HITpval 1, # confidence interval to use for HITindex significance (none, 0.75, 0.95, 0.95), HIT_CI none, # probability threshold for medium confidence with generative model [0.0, 1.0], prob_med 0.8, # probability threshold for high confidence with generative model [0.0, 1.0], prob_high 0.8.* Given the high sequencing depth in GTEx data, we raised the minimum threshold for the number of reads for confidence in HITindex classification (’—readnum’ argument) to 10 for all GTEx samples.

### PacBio Long Read Sequencing Analysis

Fastq files from PacBio reads were downloaded from the ENCODE Project data portal (https://www.encodeproject.org/)^38^, including 113 human samples from 47 tissues and cell types and 130 mouse samples from 9 tissues. cDNA-PCR PacBio sequencing data for human organoids was downloaded from the Gene Expression Omnibus (GSE203583). Long-read sequencing data was mapped to the hg38 human or mm38 reference genome using minimap2^74^ following the developers’ recommended parameters: *-ax splice:hq -uf* and *-ax splice* for PacBio and cDNA-PCR reads respectively. Reads were divided into multiple split features to define exons using *bedtools bamtobed*^75^ and assigned to overlapping genes. Only primary alignments and reads with all features assigned to the same gene were kept for downstream analyses. A detailed description of the long read sequencing samples can be found in **Supplementary Table 1**.

To retain full length reads, we then conditioned on reads whose 5’ end was within 20 nt of an empirically determined first exon identified in the full set of GTEx v8 samples^36^ for human and a subset of ENCODE samples^76^ for mouse, processed by the HITindex pipeline, and whose 3’ end was within 50nt of an experimentally verified human or mouse PAS in the polyASite 2.0 database^77^. More than 80% of PacBio reads passed these filtering criteria^40^. In addition, full unspliced reads or reads whose 5’ and 3’ end mapped to the same exon were discarded.

### TIF-seq analysis

Processed TIF-seq2 data containing the coordinates and counts of transcript start-end pairs from K562 cells was downloaded from the Gene Expression Omnibus (GSE140912). Each start and end site was intersected with all human genes and pairs which met the following criteria were retained for downstream analyses: (1) both termini aligned to the same gene, (2) the start feature was within 20 nt of an empirically derived first exon (as identified above), and (3) the end feature was within 50nt of an experimentally verified human PASs.

### Identifying PITA genes

To identify PITA genes, only genes with at least 10 reads and isoforms with at least 2 reads were considered. Genes were classified having dual alternative or solo termini by assessing the total number of first exon and polyA peaks used in each sample. Dual alternative termini genes were defined as those using at least two first exons and at least two polyA peaks, while solo termini genes were using only one first exon or polyA peak (but reads could have different specific coordinates). All mapped reads per gene per sample were used to calculate a Spearman ρ between the start and end coordinates of the reads. Genes were classified as anti-PITA, no PITA or PITA using Spearman ρ thresholds of being ρ ≤ − 0. 2, − 0. 2 > ρ < 0. 2, or ρ ≥ 0. 2 respectively.

To quantify the enrichment of PITA genes, we first calculated an area under the curve for positive and negative Spearman ρs using the *density.default* function in R to calculate AUCs using n = 512 & cut = 3. Only samples with more than 1000 genes were considered for the analysis. ΔAUCs were calculated by computing *AUC*_ρ>0_ − *AUC*_ρ<0_ .

#### Inter-tissue PITA genes

To identify inter-tissue PITA genes, we define tissue-dual alternative termini genes as those classified as solo termini in individual tissues. To avoid tissue-specific gene expression biases, we performed 100 sampling iterations, selecting two reads per isoform per gene per tissue-dual alternative termini, and then calculated the Spearman ρ between the start and end coordinates of the reads for all the sampled reads across tissues. Genes were then re-classified as solo or dual alternative termini genes based on the number of first exons and polyA peaks used across tissues.

#### Calculating genomic distances

Genomic distances were calculated using estimated coordinates for first exons and polyA peaks (as described above) for the following intervals: (1) gene length: *PAS_n_* − *TSS*_1_, (2) pre-mRNA length of PITA isoforms: *PAS*_1_ − *TSS*_1_ and *PAS_n_* − *TSS_n_*, (3) TSS interval: *TSS_n_* − *TSS*_1_, (4) internal pre-mRNA interval: *PAS*_1_ − *TSS_n_* only calculated when *TSS_n_* is upstream of *PAS*_1_, (5) PAS interval: *PAS_n_* − *PAS*_1_.

### Identifying dominant promoters with the LATER pipeline

To identify dominant promoters as defined in Alfonso-Gonzalez *et al.*^8^, the LATER pipeline (https://github.com/hilgers-lab/LATER) was applied to unfiltered human PacBio and cDNA-PCR samples following with recommended parameters^8^. Genes were classified as having promoter dominance when promoter usage was higher than 0.2 and end dominance when PAS usage was higher than 0.6^8^.

### Analysis of protein domains

Genomic coordinates for protein domains were obtained through the InterPro portal^78^. Dual alternative termini genes were overlapped with the annotated protein domains. The proportion of genes in which different isoforms overlap different protein domains was calculated for the pre-mRNA regions of isoforms with support from more than 2 reads.

### Conservation analysis

For conservation analyses, we considered the two highest expressed isoforms (most LRS reads) from the tissue with the highest Spearman ρ. Genes with Spearman ρ > 0.2 were considered to be PITA genes and all others were used as a control set. The transcript start and end coordinates from the two isoforms were ordered by genomic position and the conservation in a 400 nt genomic region centered around each site was analyzed. Conservation scores were obtained by applying the *bigWigAverageOverBed* tool from *ucsctools* to extract the mean phastCons score for each region using a hg38 phastCons (100 mammal alignment) bigwig file downloaded from the UCSC database^79,80^.

### CRISPR-a experiments

HEK293T cells were cultured in Dulbecco’s modified Eagle’s medium (DMEM) containing D-Glucose (4.5 g/L), L-Glutamine and 10% fetal bovine serum (FBS). Cells were seeded into 6-well plates and transiently transfected 16 hours later, at ∼80% confluency, using lipofectamine3000 (Invitrogen L3000001) with either UniSAM empty vector (Addgene #99866) or UniSAM vector containing sgRNA targeting LEPR AFE2 promoter region (3 replicates per condition)^48^. 48 hours post transfection, total RNA was extracted using RNeasy Mini Kit according to the manufacturer’s instructions. Reverse transcription was performed in a reaction mix containing 1 μg of total RNA, using Maxima H Minus First Strand cDNA Synthesis Kit (Thermo Scientific, K1652) according to the manufacturer’s instructions. Quantitative PCR analyses were performed with SYBR green labeling (Thermo Scientific Maxima SYBR Green/ROX qPCR Master Mix (2X), K0222) on the QuantStudio 5 Real-Time PCR System (Applied Biosystems. A28574). gRNA oligos and qPCR primers used within the study are listed in Supplementary Table 2.

### Calculation of RNAPII elongation velocities

TT-seq and mNET-seq datasets from K562 cells were downloaded from the Gene Expression Omnibus (GSE148433 and GSE159633, respectively). For both TT-seq and mNET-seq datasets, initial read trimming was performed using cutadapt^81^ with parameters *--minimum-length 25*, *--quality-cutoff 25*, and *--overlap 12*. The adapter sequences -a AGATCGGAAGAGCACACGTCTGAACTCCAGTCA and -A AGATCGGAAGAGCGTCGTGTAGGGAAAGAGTGT were removed from TT-seq reads and -a TGGAATTCTCGGGTGCCAAGGAACTCCAGTCAC and -A AGATCGTCGGACTGTAGAACTCTGAAC were removed from mNET-seq reads. Both trimmed datasets were then aligned to the human reference genome GRCh38.v43 using STAR^73^, with alignment parameters*--quantMode GeneCounts*, *--outSAMtype BAM SortedByCoordinate*, *--outFilterType BySJout*, *--outFilterMultimapNmax 1*, *--alignSJoverhangMin 8*, *--alignSJDBoverhangMin 1*, *--outFilterMismatchNmax 999*, *--outFilterMismatchNoverLmax 0.02*, *--alignIntronMin 20*, *--alignIntronMax 1000000*, and *--alignMatesGapMax 1000000*. Spike-in reads were aligned to the yeast reference genome S.cerevisiae.R64 and ERCC synthetic spiked-in sequences as described in^82^, for the purpose of normalization of TT-seq counts and mNET-seq counts, respectively. RNAPII elongation velocities were calculated by dividing the amount of transcribed RNA (TT-seq coverage) by the density of transcribing RNAPII (mNET-seq reads), as described previously^54,55^. For each gene, the mean elongation velocity across replicates was used for downstream analyses.

### Analyses of Micro-C and epigenetics data

Tier 1 micro-C ultradeep datasets for HFFc6 4DNFI9FVHJZQ and H1-ESC 4DNFI9GMP2J8 were downloaded from the 4DN Portal^49,83^ as multi-resolution cooler files^84^ and used for analysis of interaction landscapes around PITA and no PITA genes. Using HFFc6 and H1-ESCs PacBio data, we selected the most upstream and downstream polyA and FE peak coordinates for each gene. To ensure all selected sites could be clearly visualized on micro-C heatmaps at 500bp resolution, we only used PITA and non-PITA genes with a minimum distance of 2kb between all selected sites. Average aggregated interaction profiles around genes (rescaled meta-gene pileups) were generated using coolpup.py^85^ as follows: regions of interest were extended 100% up- and down-stream (flanking) and corresponding observed/expected matrices were extracted from microC (cooler-files) at 500 bp resolution, rescaled to 200*200 pixels, matrices corresponding to negatively stranded genes were “flipped”, and averaged. Insulation profiles were calculated genome-wide for both microC datasets at 250 bp resolution using insulation window size of 2500 bp using cooltools^86^ and stored as bigwig files. Stackups of relevant epigenetic features/tracks (insulation, ATAC-seq, H3K4me3 ChIP-seq, CTCF ChIP-seq^50^) were generated by extracting bigwig signal binned into 200 bins for each genomic interval (TSS_1,n_, PAS_1,n_ flanked with 7,500 bp upstream and downstream) using pyBBI package^87^ (https://zenodo.org/records/10382981). Insulation profiles for each genomic interval were normalized by subtracting mean flanking signal (50 upstream most and 50 downstream most bins) as in ^52^.

## CODE AVAILABILITY

Code for analyses of short and long-read sequencing datasets is available at https://github.com/thepailab/mrna_initiation_termination. Code for microC-related analyses is available at https://github.com/dekkerlab/pita_project.

## Acknowledgements

We thank Zachary Wakefield for providing a curated list of human transcription start sites, Marian Walhout for suggesting the interspecies comparative analyses, Phil Zamore for discussions, and Fiszbein and Pai labs for helpful discussions and comments. This work was funded by grants from the National Institutes of Health (R35GM133762 and R01HG012967 to A.A.P.; R35GM147254 to A.F.; and R01HG003143 and UM1HG011536 to J.D.). J.D. is an investigator of the Howard Hughes Medical Institute.

## Author Contributions

Contributed equally (ECR, CLC), Conceived and designed experiments (AF, AAP), Performed the experiments (ECR, CLC), Performed statistical analyses (ECR, CLC, SVV), Analyzed the data (ECR, CLC, SVV, GYK, SM, AF, AAP), Contributed reagents/materials/analysis tools (JD, AF, AAP), Wrote the paper (AF, AAP), Jointly supervised research (AF, AAP)

## Notes

### Competing Interest Statement

The authors have declared no competing interest.

## REFERENCES

1. Wang, E. T. et al. Alternative isoform regulation in human tissue transcriptomes. Nature 456, 470–476 (2008).

2. Joglekar, A. et al. Single-cell long-read mRNA isoform regulation is pervasive across mammalian brain regions, cell types, and development. bioRxiv (2023) doi:10.1101/2023.04.02.535281.

3. Nilsen, T. W. & Graveley, B. R. Expansion of the eukaryotic proteome by alternative splicing. Nature 463, 457–463 (2010).

4. Tian, B. & Manley, J. L. Alternative polyadenylation of mRNA precursors. (2016) doi:10.1038/nrm.2016.116.

5. Anvar, S. Y. et al. Full-length mRNA sequencing uncovers a widespread coupling between transcription initiation and mRNA processing. Genome Biol. 19, 46 (2018).

6. Fiszbein, A., Krick, K. S., Begg, B. E. & Burge, C. B. Exon-Mediated Activation of Transcription Starts. Cell 179, 1551–1565.e17 (2019).

7. Hardwick, S. A. et al. Single-nuclei isoform RNA sequencing unlocks barcoded exon connectivity in frozen brain tissue. Nat. Biotechnol. 40, 1082–1092 (2022).

8. Alfonso-Gonzalez, C. et al. Sites of transcription initiation drive mRNA isoform selection. Cell 186, 2438–2455.e22 (2023).

9. Kaida, D. et al. U1 snRNP protects pre-mRNAs from premature cleavage and polyadenylation. Nature 468, 664–668 (2010).

10. Laitem, C. et al. CDK9 inhibitors define elongation checkpoints at both ends of RNA polymerase II-transcribed genes. Nat. Struct. Mol. Biol. 22, 396–403 (2015).

11. Caizzi, L. et al. Efficient RNA polymerase II pause release requires U2 snRNP function Efficient RNA polymerase II pause release requires U2 snRNP function. 1–15 (2021).

12. Nagaike, T. et al. Transcriptional activators enhance polyadenylation of mRNA precursors. Mol. Cell 41, (2011).

13. Guo, Y. E. et al. Pol II phosphorylation regulates a switch between transcriptional and splicing condensates. Nature 572, 543–548 (2019).

14. Shabalina, S. A., Ogurtsov, A. Y., Spiridonov, N. A. & Koonin, E. V. Evolution at protein ends: Major contribution of alternative transcription initiation and termination to the transcriptome and proteome diversity in mammals. Nucleic Acids Res. 42, 7132–7144 (2014).

15. A promoter-level mammalian expression atlas. Nature 507, 462–470 (2014).

16. Stroup, E. K. & Ji, Z. Deep learning of human polyadenylation sites at nucleotide resolution reveals molecular determinants of site usage and relevance in disease. Nat. Commun. 14, 7378 (2023).

17. Taliaferro, J. M. et al. Distal Alternative Last Exons Localize mRNAs to Neural Projections. Mol. Cell 61, 821–833 (2016).

18. Lianoglou, S., Garg, V., Yang, J. L., Leslie, C. S. & Mayr, C. Ubiquitously transcribed genes use alternative polyadenylation to achieve tissue-specific expression. Genes and Development 27, 2380–2396 (2013).

19. Di Giammartino, D. C., Nishida, K. & Manley, J. L. Mechanisms and Consequences of Alternative Polyadenylation. Molecular Cell vol. 43 853–866 Preprint at 10.1016/j.molcel.2011.08.017 (2011).

20. Pal, S. et al. Alternative transcription exceeds alternative splicing in generating the transcriptome diversity of cerebellar development. Genome Res. 21, 1260–1272 (2011).

21. Pai, A. A. & Luca, F. Environmental influences on RNA processing: Biochemical, molecular and genetic regulators of cellular response. Wiley Interdiscip. Rev. RNA (2019).

22. Reyes, A. & Huber, W. Alternative start and termination sites of transcription drive most transcript isoform differences across human tissues. Nucleic Acids Res. 46, 582–592 (2018).

23. Pabitra K. Parua Sampada Kalan Bradley Benjamin, M. S. &. R. P. F. Distinct Cdk9-phosphatase switches act at the beginning and end of elongation by RNA polymerase II. Nat. Commun. (2020) doi:10.1038/s41467-020-18173-6.

24. Parua, P. K. et al. A Cdk9-PP1 switch regulates the elongation-termination transition of RNA polymerase II. Nature 558, 460–464 (2018).

25. Kaplan, C. D., Jin, H., Zhang, I. L. & Belyanin, A. Dissection of Pol II trigger loop function and Pol II activity-dependent control of start site selection in vivo. PLoS Genet. 8, e1002627 (2012).

26. Braberg, H. et al. From structure to systems: high-resolution, quantitative genetic analysis of RNA polymerase II. Cell 154, 775–788 (2013).

27. Qiu, C. et al. Universal promoter scanning by Pol II during transcription initiation in Saccharomyces cerevisiae. Genome Biol. 21, 132 (2020).

28. Geisberg, J. V., Moqtaderi, Z. & Struhl, K. The transcriptional elongation rate regulates alternative polyadenylation in yeast. Elife 9, (2020).

29. Geisberg, J. V. et al. Nucleotide-level linkage of transcriptional elongation and polyadenylation. Elife 11, (2022).

30. Cortazar, M. A. et al. Control of RNA Pol II Speed by PNUTS-PP1 and Spt5 Dephosphorylation Facilitates Termination by a ‘Sitting Duck Torpedo’ Mechanism. Mol. Cell 76, 896–908.e4 (2019).

31. Goering, R. et al. LABRAT reveals association of alternative polyadenylation with transcript localization, RNA binding protein expression, transcription speed, and cancer survival. doi:10.1101/2020.10.05.326702.

32. Rowley, M. J. et al. Condensin II Counteracts Cohesin and RNA Polymerase II in the Establishment of 3D Chromatin Organization. Cell Rep. 26, 2890–2903.e3 (2019).

33. Wu, H., Zhang, J., Tan, L. & Sunney Xie, X. Extruding transcription elongation loops observed in high-resolution single-cell 3D genomes. bioRxiv 2023.02.18.529096 (2023) doi:10.1101/2023.02.18.529096.

34. Leidescher, S. et al. Spatial organization of transcribed eukaryotic genes. Nat. Cell Biol. 24, 327–339 (2022).

35. Fiszbein, A. et al. Widespread occurrence of hybrid internal-terminal exons in human transcriptomes. Sci Adv 8, eabk1752 (2022).

36. GTEx Consortium. The GTEx Consortium atlas of genetic regulatory effects across human tissues. Science 369, 1318–1330 (2020).

37. Katz, Y., Wang, E. T., Airoldi, E. M. & Burge, C. B. Analysis and design of RNA sequencing experiments for identifying isoform regulation. Nat. Methods 7, 1009–1015 (2010).

38. Luo, Y. et al. New developments on the Encyclopedia of DNA Elements (ENCODE) data portal. Nucleic Acids Res. 48, D882–D889 (2020).

39. Reese, F., et al. The ENCODE4 long-read RNA-seq collection reveals distinct classes of transcript structure diversity. bioRxiv (2023) doi:10.1101/2023.05.15.540865.

40. Calvo-Roitberg, E., Daniels, R. F. & Pai, A. A. Challenges in identifying mRNA transcript starts and ends from long-read sequencing data. bioRxiv (2023) doi:10.1101/2023.07.26.550536.

41. Dong, X. et al. Benchmarking long-read RNA-sequencing analysis tools using in silico mixtures. Nat. Methods 20, 1810–1821 (2023).

42. Mayr, C. & Bartel, D. P. Widespread shortening of 3’UTRs by alternative cleavage and polyadenylation activates oncogenes in cancer cells. Cell 138, 673–684 (2009).

43. Sandberg, R., Neilson, J. R., Sarma, A., Sharp, P. A. & Burge, C. B. Proliferating cells express mRNAs with shortened 3’ untranslated regions and fewer microRNA target sites. Science 320, 1643–1647 (2008).

44. Ji, Z., Lee, J. Y., Pan, Z., Jiang, B. & Tian, B. Progressive lengthening of 3’ untranslated regions of mRNAs by alternative polyadenylation during mouse embryonic development. Proc. Natl. Acad. Sci. U. S. A. 106, 7028–7033 (2009).

45. Hou, R., Hon, C.-C. & Huang, Y. CamoTSS: analysis of alternative transcription start sites for cellular phenotypes and regulatory patterns from 5’ scRNA-seq data. bioRxiv (2023) doi:10.1101/2023.04.17.536840.

46. Richards, A. L. et al. Environmental perturbations lead to extensive directional shifts in RNA processing. PLoS Genet. 13, e1006995 (2017).

47. Tajnik, M. et al. Intergenic Alu exonisation facilitates the evolution of tissue-specific transcript ends. Nucleic Acids Res. 43, 10492–10505 (2015).

48. Fidanza, A. et al. An all-in-one UniSam vector system for efficient gene activation. Sci. Rep. 7, 6394 (2017).

49. Krietenstein, N. et al. Ultrastructural Details of Mammalian Chromosome Architecture. Mol. Cell 78, 554–565.e7 (2020).

50. Akgol Oksuz, B., et al. Systematic evaluation of chromosome conformation capture assays. Nat. Methods 18, 1046–1055 (2021).

51. Bonev, B. et al. Multiscale 3D Genome Rewiring during Mouse Neural Development. Cell 171, 557–572.e24 (2017).

52. Valton, A.-L. et al. A cohesin traffic pattern genetically linked to gene regulation. Nat. Struct. Mol. Biol. 29, 1239–1251 (2022).

53. Schwalb, B. et al. TT-seq maps the human transient transcriptome. Science 352, 1225–1228 (2016).

54. Žumer, K., et al. Two distinct mechanisms of RNA polymerase II elongation stimulation in vivo. Mol. Cell 81, 3096–3109.e8 (2021).

55. Caizzi, L. et al. Efficient RNA polymerase II pause release requires U2 snRNP function. Mol. Cell 81, 1920–1934.e9 (2021).

56. Veloso, A. et al. Rate of elongation by RNA polymerase II is associated with specific gene features and epigenetic modifications. Genome Res. 24, 896–905 (2014).

57. Fong, N. et al. Pre-mRNA splicing is facilitated by an optimal RNA polymerase II elongation rate. Genes Dev. 28, 2663–2676 (2014).

58. Maslon, M. M. et al. A slow transcription rate causes embryonic lethality and perturbs kinetic coupling of neuronal genes. EMBO J. 38, (2019).

59. Zhang, Z., Bae, B., Cuddleston, W. H. & Miura, P. Coordination of alternative splicing and alternative polyadenylation revealed by targeted long read sequencing. Nat. Commun. 14, 5506 (2023).

60. Glover-Cutter, K., Kim, S., Espinosa, J. & Bentley, D. L. RNA polymerase II pauses and associates with pre-mRNA processing factors at both ends of genes. Nat. Struct. Mol. Biol. 15, 71–78 (2008).

61. Takahashi, H. et al. The role of Mediator and Little Elongation Complex in transcription termination. Nat. Commun. 11, 1063 (2020).

62. Ullah, F., Jabeen, S., Salton, M., Reddy, A. S. N. & Ben-Hur, A. Evidence for the role of transcription factors in the co-transcriptional regulation of intron retention. Genome Biol. 24, 53 (2023).

63. Thompson, M. et al. Splicing in a single neuron is coordinately controlled by RNA binding proteins and transcription factors. (2019) doi:10.7554/eLife.46726.

64. Kwek, K. Y. et al. U1 snRNA associates with TFIIH and regulates transcriptional initiation. Nat. Struct. Biol. 9, 800–805 (2002).

65. Uriostegui-Arcos, M., Mick, S. T., Shi, Z., Rahman, R. & Fiszbein, A. Splicing activates transcription from weak promoters upstream of alternative exons. Nat. Commun. 14, 3435 (2023).

66. Bergfort, A. & Neugebauer, K. M. The promoter as a trip navigator: Guiding alternative polyadenylation site destinations. Molecular cell vol. 83 2395–2397 (2023).

67. Ibrahim, M. M. et al. Determinants of promoter and enhancer transcription directionality in metazoans. Nat. Commun. 9, 4472 (2018).

68. Han, L. et al. Concentration and length dependence of DNA looping in transcriptional regulation. PLoS One 4, e5621 (2009).

69. Chen, L. et al. R-ChIP Using Inactive RNase H Reveals Dynamic Coupling of R-loops with Transcriptional Pausing at Gene Promoters. Mol. Cell 68, 745–757.e5 (2017).

70. Moabbi, A. M., Agarwal, N., El Kaderi, B. & Ansari, A. Role for gene looping in intron-mediated enhancement of transcription. Proc. Natl. Acad. Sci. U. S. A. 109, 8505–8510 (2012).

71. Agarwal, N. & Ansari, A. Enhancement of Transcription by a Splicing-Competent Intron Is Dependent on Promoter Directionality. PLoS Genet. 12, e1006047 (2016).

72. Kwon, B. et al. Enhancers regulate 3’ end processing activity to control expression of alternative 3’UTR isoforms. Nat. Commun. 13, 2709 (2022).

73. Dobin, A. et al. STAR: ultrafast universal RNA-seq aligner. Bioinformatics 29, 15–21 (2013).

74. Li, H. New strategies to improve minimap2 alignment accuracy. Bioinformatics 37, 4572–4574 (2021).

75. Quinlan, A. R. & Hall, I. M. BEDTools: A flexible suite of utilities for comparing genomic features. Bioinformatics 26, 841–842 (2010).

76. Lin, S. et al. Comparison of the transcriptional landscapes between human and mouse tissues. Proc. Natl. Acad. Sci. U. S. A. 111, 17224–17229 (2014).

77. Herrmann, C. J. et al. PolyASite 2.0: a consolidated atlas of polyadenylation sites from 3′ end sequencing. Nucleic Acids Res. 48, D174–D179 (2019).

78. Paysan-Lafosse, T. et al. InterPro in 2022. Nucleic Acids Res. 51, D418–D427 (2023).

79. Siepel, A. et al. Evolutionarily conserved elements in vertebrate, insect, worm, and yeast genomes. Genome Res. 15, 1034–1050 (2005).

80. Raney, B. J. et al. The UCSC Genome Browser database: 2024 update. Nucleic Acids Res. (2023) doi:10.1093/nar/gkad987.

81. Martin, M. Cutadapt removes adapter sequences from high-throughput sequencing reads. EMBnet.journal 17, 10–12 (2011).

82. Wachutka, L., Caizzi, L., Gagneur, J. & Cramer, P. Global donor and acceptor splicing site kinetics in human cells. Elife 8, (2019).

83. Reiff, S. B. et al. The 4D Nucleome Data Portal as a resource for searching and visualizing curated nucleomics data. Nat. Commun. 13, 2365 (2022).

84. Abdennur, N. & Mirny, L. A. Cooler: scalable storage for Hi-C data and other genomically labeled arrays. Bioinformatics 36, 311–316 (2020).

85. Flyamer, I. M., Illingworth, R. S. & Bickmore, W. A. Coolpup.py: versatile pile-up analysis of Hi-C data. Bioinformatics 36, 2980–2985 (2020).

86. Open2C et al. Cooltools: enabling high-resolution Hi-C analysis in Python. bioRxiv 2022.10.31.514564 (2022) doi:10.1101/2022.10.31.514564.

87. Kent, W. J., Zweig, A. S., Barber, G., Hinrichs, A. S. & Karolchik, D. BigWig and BigBed: enabling browsing of large distributed datasets. Bioinformatics 26, 2204–2207 (2010).

88. Wang, J. et al. TIF-Seq2 disentangles overlapping isoforms in complex human transcriptomes. Nucleic Acids Res. 48, e104 (2020).

