## Supplementary Figures for "mRNA initiation and termination are spatially coordinated"

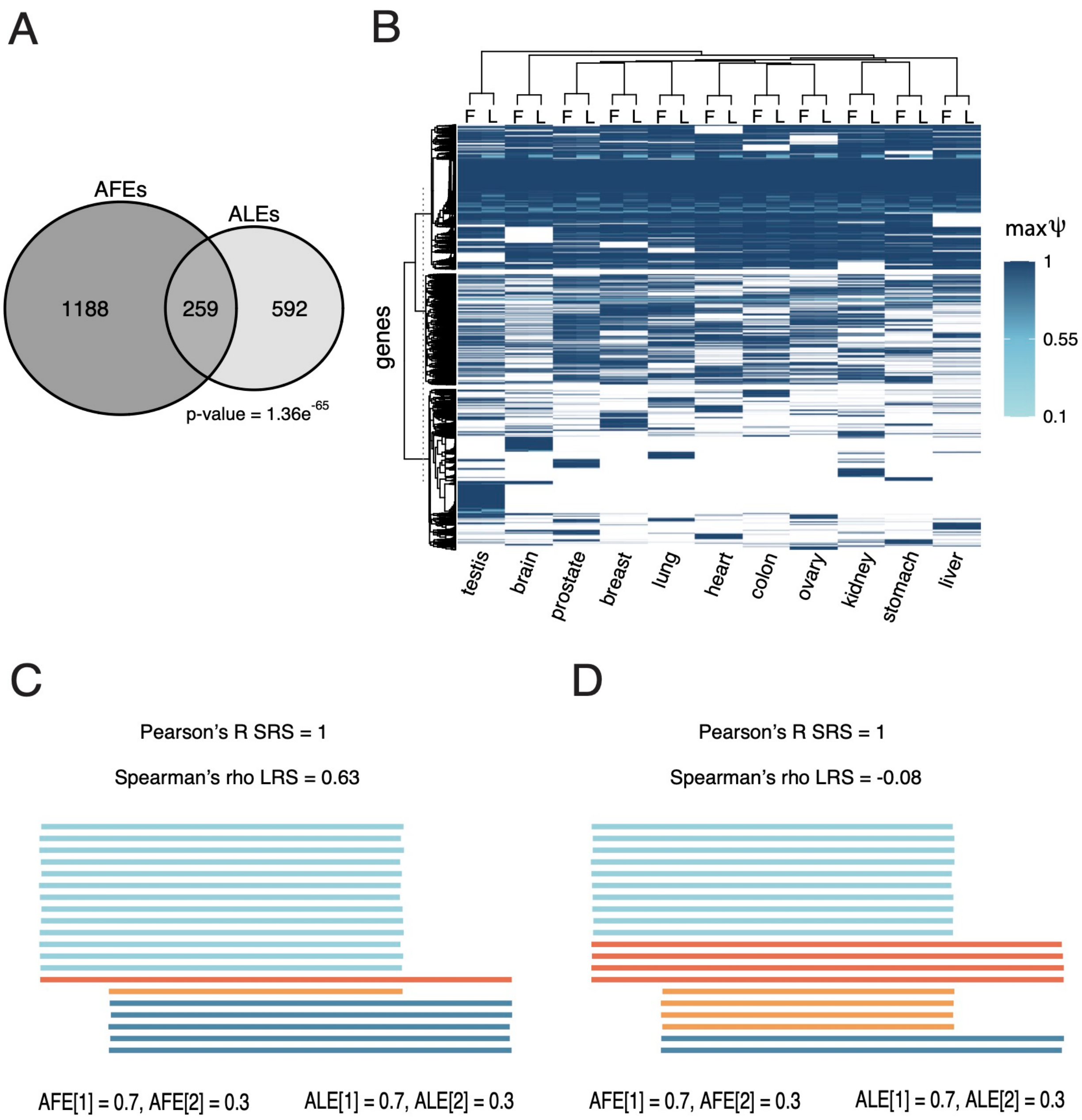


**Extended Data Figure 1: Assessing frequency of alternative first and last exon usage. (A).** Overlap in genes with both AFEs and ALEs (mean across all samples). p-value = $1.36 \times 10^{-65}$. **(B)** Maximum PSI (Ψ) value of alternative first (*F*) and last exons *(L)* per gene (*rows*) across 11 tissues *(columns*). White indicates no gene expression in the given tissue. Dendrogram indicates hierarchical clustering of genes, showing that max Ψ values for AFE and ALEs tend to be similar within a gene. Schematics of a gene with direct **(C)** and indirect **(D)** correlations between first and last exon usage, where each line represents a hypothetical read and Spearman's ρ values are indicated. In both instances, AFE and ALE usage is correlated based on ordinal position and short-read sequencing analyses would result in a Pearson's R = 1.

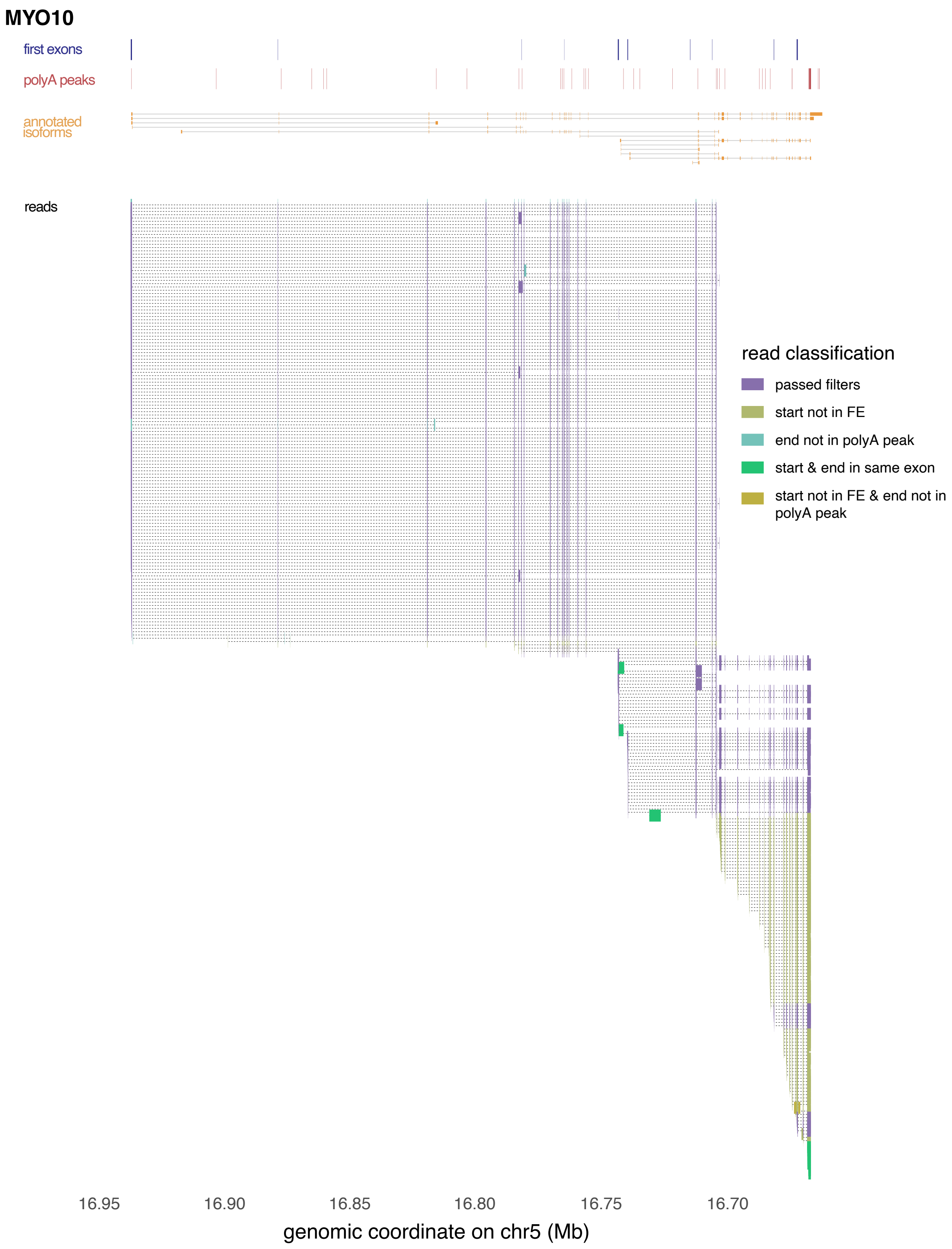


**Extended Data Figure 2: Filtering long read sequencing data.** HITindex first exons (*blue, first track*), polyA-seq peaks (*red, second track*), annotated isoforms (*orange, third track*), and all unfiltered LRS reads for *MYO10* (introns in dotted black lines). Reads are colored based on meeting criteria to remove reads with spurious start and end positions.

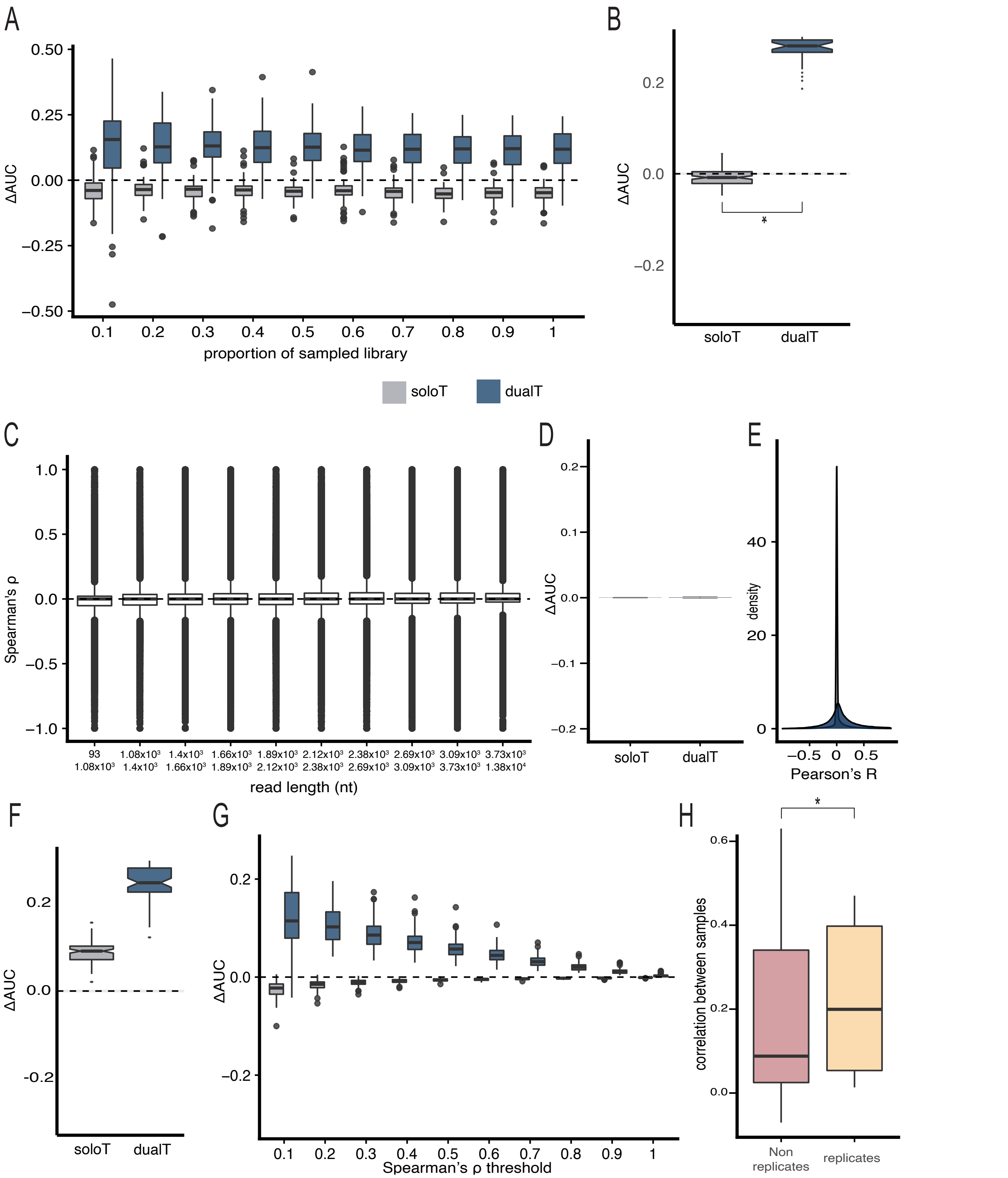


**Extended Data Figure 3**: **Assessing biases in identification of PITA genes.** (**A**) Distributions of ΔAUC values across tissues (*y-axis*) for varying subsampled libraries with varying read depth (*x-axis*). (**B**) Distribution of ΔAUC values across tissues (y-axis) calculated after sampling two reads for each expressed isoform per dual alternative and solo termini gene. (**C**) Distribution of Spearman ρs (*y-axis*) across deciles of mean read lengths per gene (*x-axis*). (**D**) Distribution of ΔAUCs per sample (y-axis) for solo and dual alternative termini genes calculated after permuting relationship between read starts and ends within each sample. T-test * p-value < $10^{-16}$. Distribution of Pearson's R across genes (E) and ΔAUCs (F) when evaluating the direct relationship between the ordinal position of first exons and polyA peaks in long read sequencing data. (**G**) Distribution of ΔAUCs (*y-axis*) across varying Spearman ρ thresholds x (*x-axis*), where $\Delta AUC = AUC_{R>x} - AUC_{R<-x}$ . (**H**) Distribution of Pearson Correlations (y-axis) between Spearman's ρ distributions between samples that are replicates and non-replicates. Wilcoxon test * p-value < $10^{-4}$.

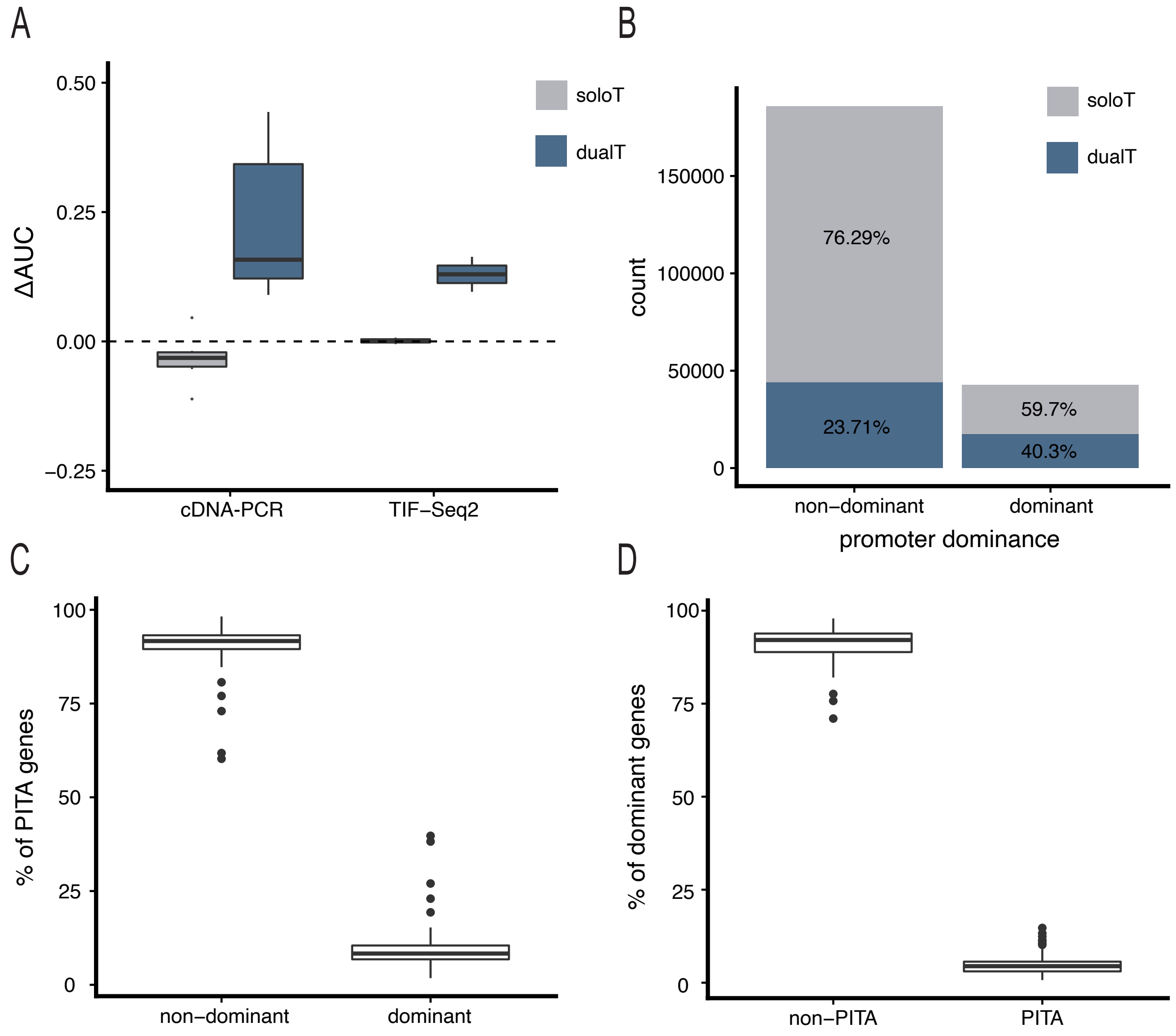


**Extended Data Figure 4**: **PITA is consistent across sequencing techniques.** (**A**) Distribution of ΔAUCs for cDNA-PCR long read sequencing data [8] and TIF-Seq2 short-read sequencing data [88]. (**B**) Number of genes classified as having non-dominant or dominant promoters in the human Iso-seq data after analysis with the LATER pipeline (CITE). The counts of genes that have solo or dual alternative termini are indicated in grey and blue, respectively. (**C**) Percentage of PITA genes in Iso-seq data that are classified as having non-dominant or dominant promoter by the LATER pipeline. (**D**) Percentage of genes with dominant promoters in Iso-seq data that are classified as non-PITA or PITA.

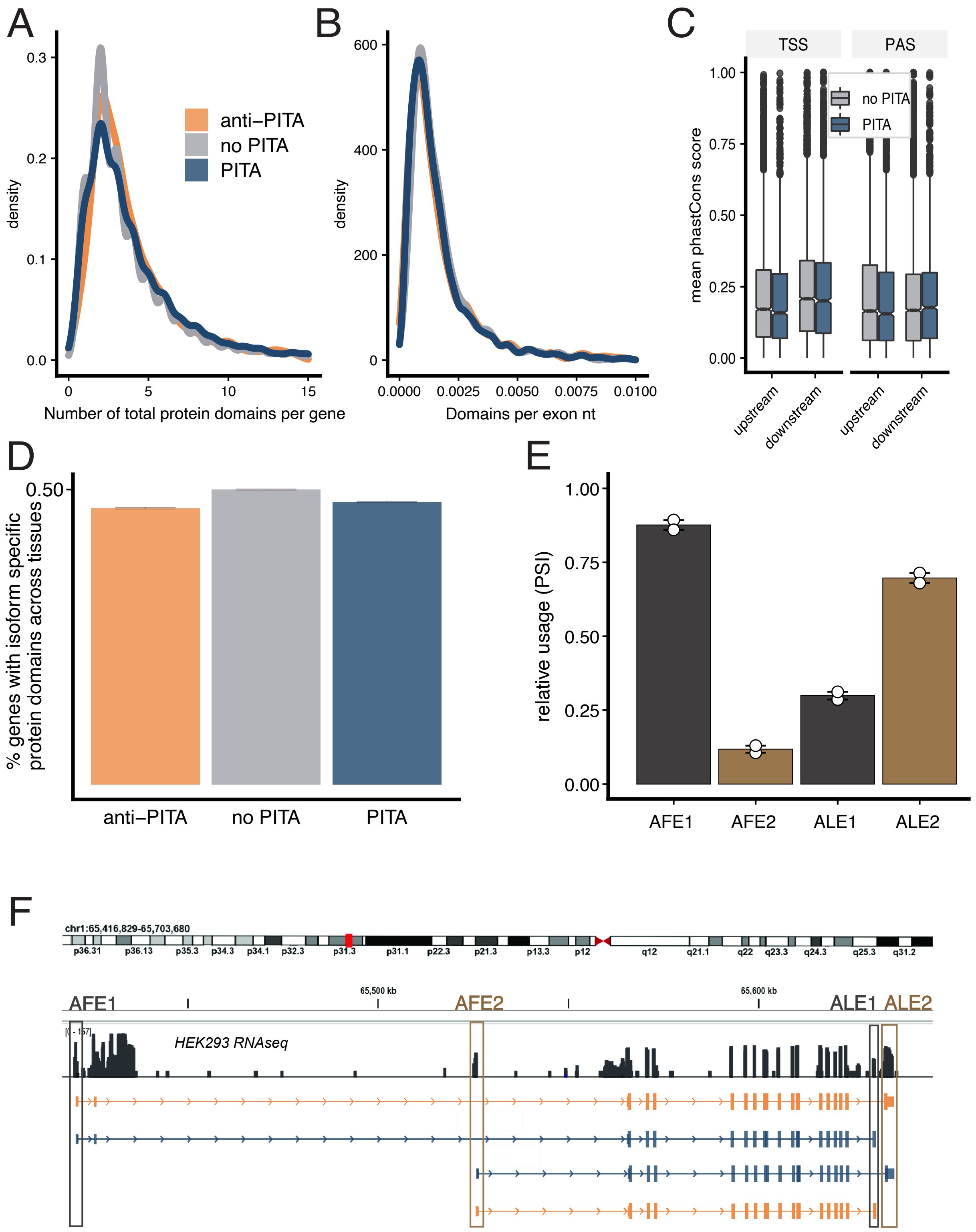


**Extended Data Figure 5: Properties of PITA genes.**(**A**) Distribution of the number of annotated protein domains per gene for different classes of genes. (**B**) Distribution of the number of annotated protein domainsper gene normalized by mRNA length across different classes of genes. (**C**) Conservation scores (mean phastCons score, *y-axis*) in a 400nt region around each terminal site of the two most highly expressed isoforms for tissue dual alternative termini genes with dual alternative termini. (**D**) Proportion of tissue dual alternative termini genes (*y-axis*) whose isoforms overlap different annotated protein domains. **(E)** Mean PSI values in HEK293 cells for each terminal exon in *LEPR*. **(F)** Raw short-read RNA-seq coverage in HEK293 cells (*top*) and annotated isoform (*bottom*) for *LEPR*.

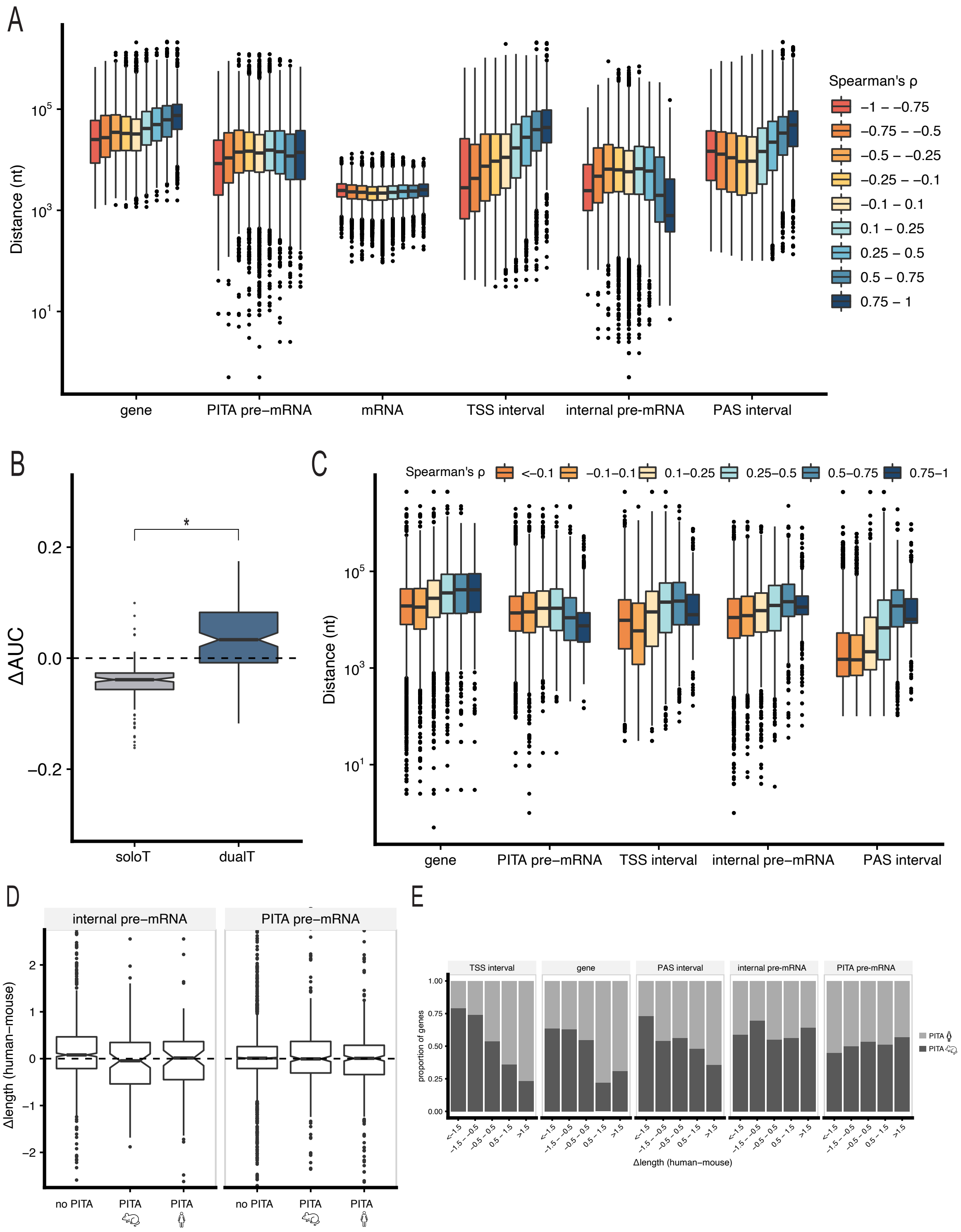


**Extended Data Figure 6: Relationship between gene architecture and PITA.**(**A**) Distributions of genomic distances (*y-axis*) across bins of Spearman's ρ for dual alternative termini genes for gene, PITA pre-mRNA, average mRNA, TSS, internal pre-mRNA and PAS intervals for human genes. (**B**) Distribution of ΔAUC values across 130 ENCODE mouse long-read sequencing samples from 9 tissue and cell types for genes using solo (*gray*) or dual alternative termini (*blue*), T-test * p-value $< 10^{-16}$. (**C**) Distributions of genomic distances (*y-axis*) across bins of Spearman's ρ for values of dual alternative termini genes for gene,, PITA pre-mRNAs, average mRNA, TSS, internal pre-mRNA and between TSS the maximum distances between TSSs, the downstream-most TSS and upstream-most PAS (internal pre-mRNA), and PAS intervals for mouse genes. (**D**) Distribution of the difference in internal pre-mRNA and PITA pre-mRNA lengths for human and mouse orthologs (*y-axis*) of genes that are not PITA in either species, PITA only in mice or PITA only in humans. To account for global differences in gene lengths between species, distances were first normalized by the mean distance within each species for each feature. No significant differences were observed. **(E)** Proportion of genes (*y-axis*) that are only PITA in human (*light grey*) or only PITA in mouse (*dark grey*) for different Δlength bins across gene lengths, PITA pre-mRNA lengths, TSS intervals, internal pre-mRNA intervals, or PAS intervals.

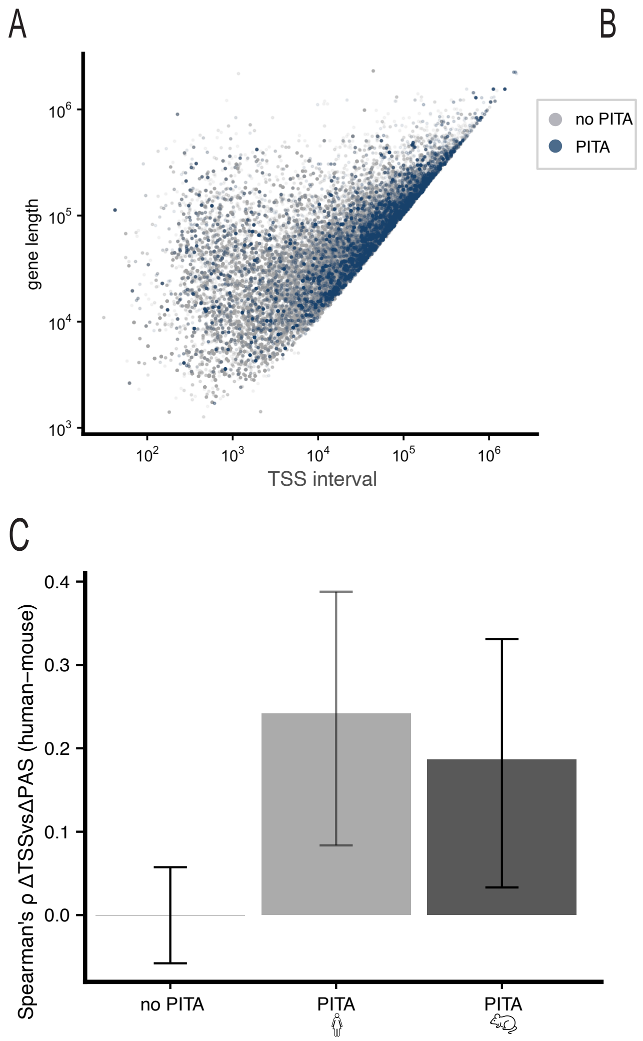


**Extended Data Figure 7. Relationships between TSS and PAS intervals across genes.** (**A**) Correlation between gene length (*y-axis*) and the TSS interval (*x-axis*) for non PITA and PITA genes (Pearson's R = 0.70 and 0.89, respectively). (**B**) Correlation between gene length (*y-axis*) and PAS interval for non PITA and PITA genes (Pearson's R = 0.71 and 0.84, respectively). (**C**) Mean correlation between the ΔTSS interval and ΔPAS interval (*y-axis*) for no PITA, mouse-specific PITA, and human-specific PITA genes. Error bars indicate 95% confidence intervals.

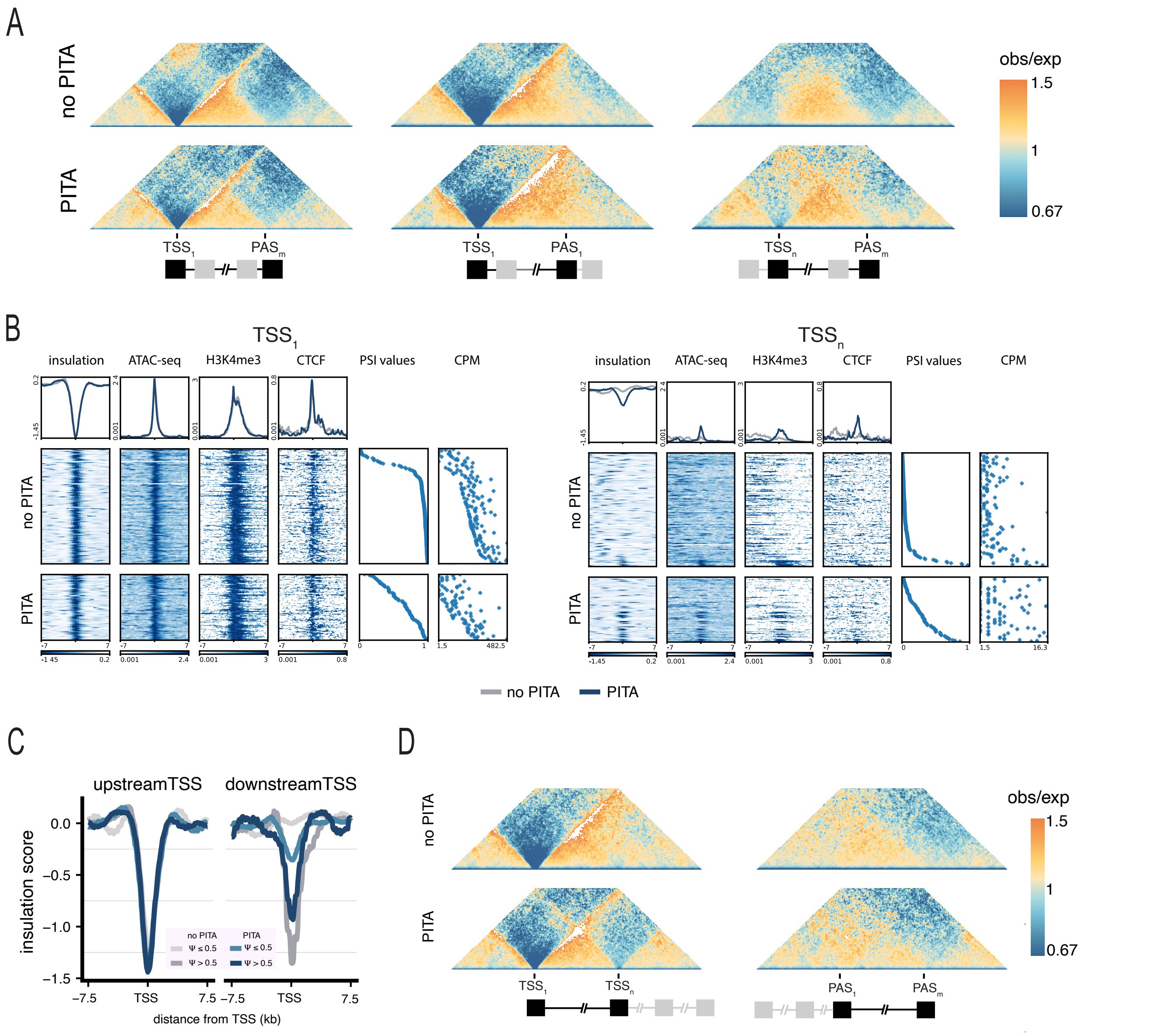


**Extended Data Fig 8. microC maps in Human Foreskin Fibroflast cells. (A)** Aggregated and scaled micro-C maps for TSS1-PASn, TSS1-PAS1, and TSSn-PASm pairs at 500 bp resolution in HFFs, where the color scale indicates low (depletion of interactions, *blue*) to high (enrichment of interactions, *orange*) microC signal enrichment, where each figure pixel reflects a paired interaction. **(B)** Stackups of normalized insulation scores, ATAC-seq signal, H3K4me3 ChIP-seq signal, and CTCF ChIP-seq signal for active TSSs in non-PITA and PITA genes in HFFs, sorted based on relative site usage (PSI values). Also shown are PSI values and counts per million (CPMs). Stackups are flipped according to the orientation of the genes, to have the gene body on the right of the TSSs. **(C)** Average insulation profiles in HFFs at 250 bp resolution in the +/-7.5kb window around upstream $TSS_1$ (*left*) and downstream $TSS_n$ (*right*) for non-PITA (*grey scale*) and PITA (*blue scale*) genes. Shades indicate relative TSS usage (measured by PSI values). **(D)** Aggregated and scaled micro-C maps for TSS1-TSSn, PAS1-PASm pairs at 500 bp resolution in HFFs, where the color scale indicates high (*orange*) to low (*blue*)microC signal enrichment.

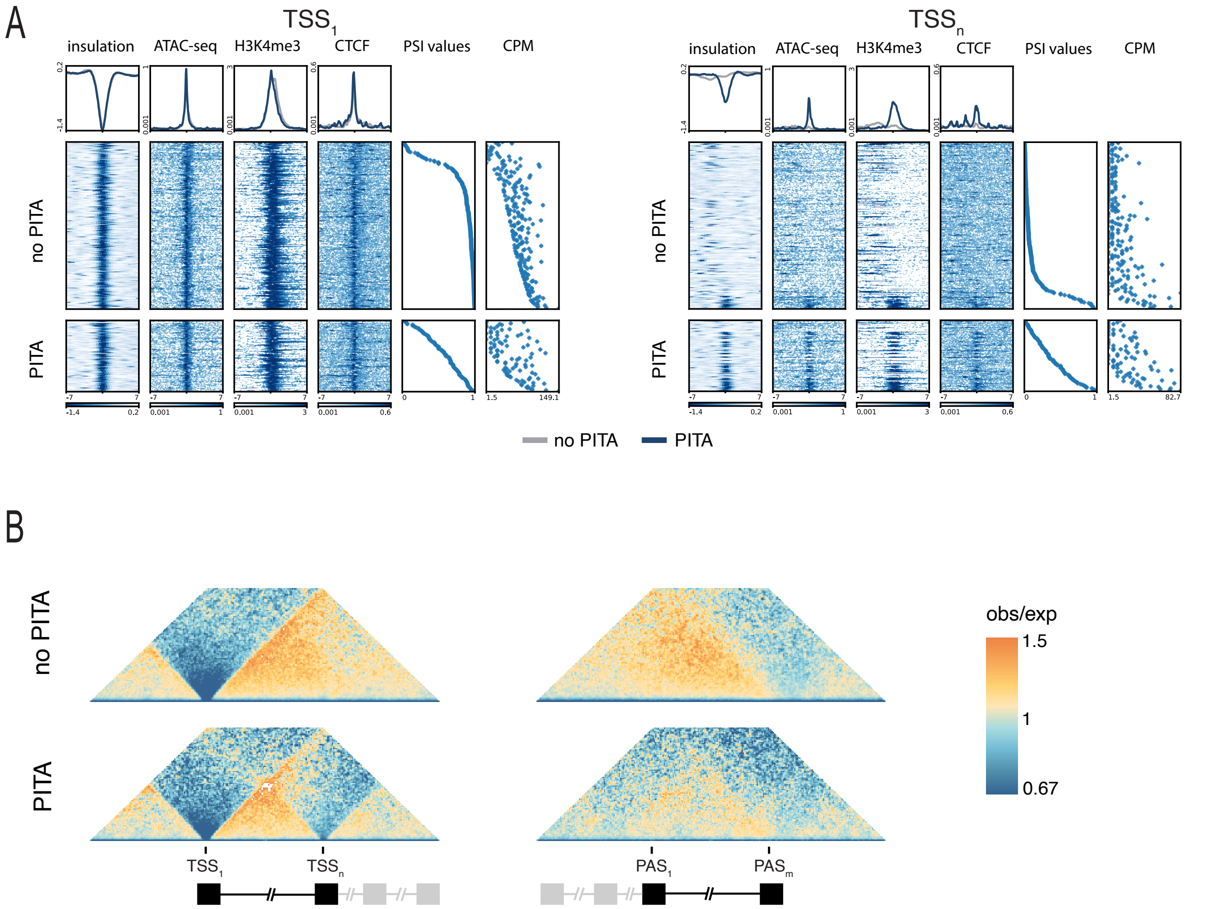


**Extended Data Figure 9. microC maps in Human 1 embryonic stem cells. (A)** Stackups of normalized insulation scores, ATAC-seq signal, H3K4me3 ChIP-seq signal, and CTCF ChIP-seq signal for active TSSs in non-PITA and PITA genes in H1-ESCs, sorted based on relative site usage (PSI values). Also shown are PSI values and counts per million (CPMs). Stackups are flipped according to the orientation of the genes, to have the gene body on the right of the TSSs. **(B)** Aggregated and scaled micro-C maps for TSS1-TSSn, PAS1-PASm pairs at 500 bp resolution in H1-ESCs, where the color scale indicates high (*orange*) to low (*blue*) microC signal enrichment.

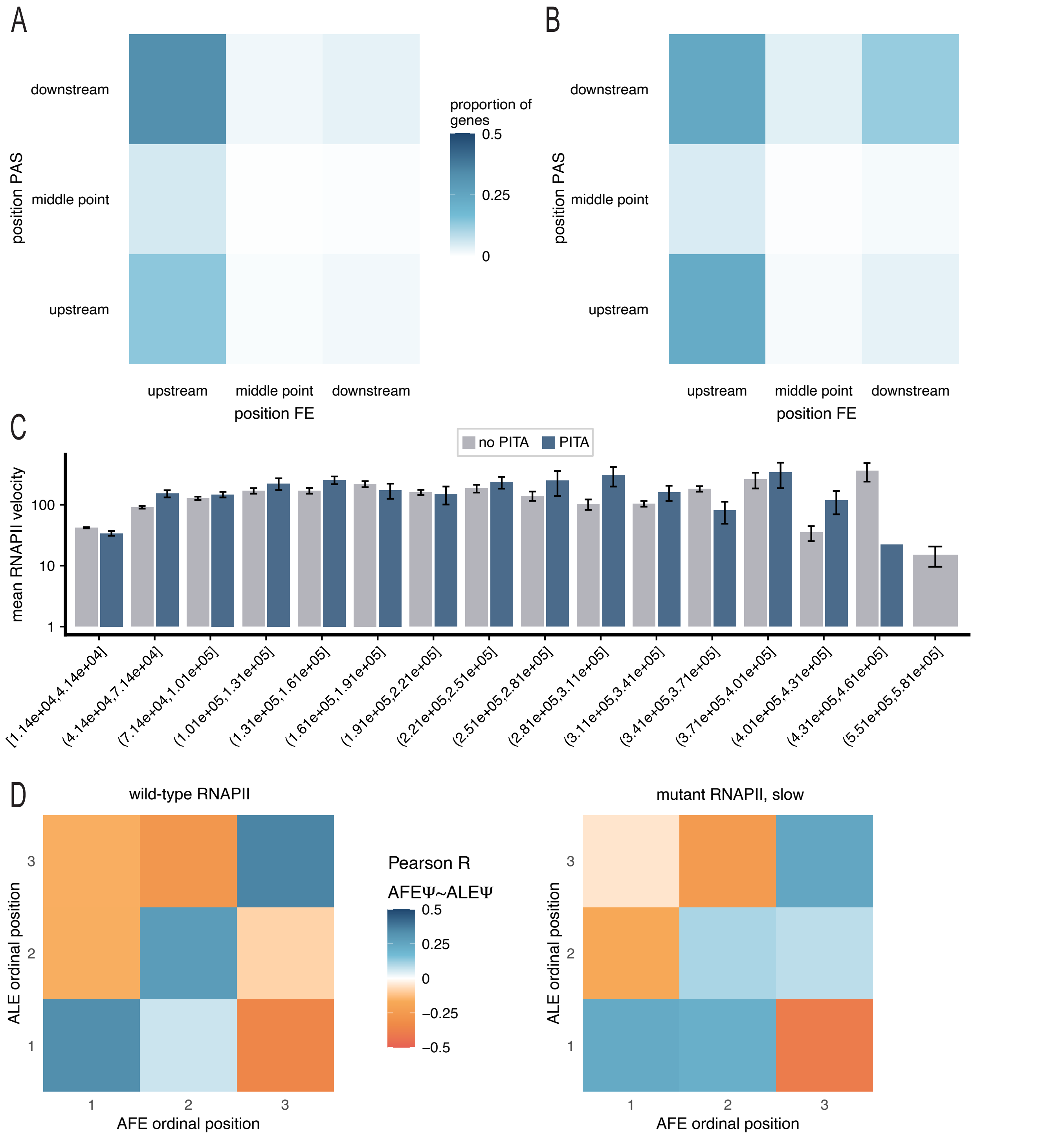


**Extended Data Figure 10. Relationship between PITA and elongation rates.** Proportion of reads per sample (*y-axis*) that use a given FE position (*x-axis*) and a given PAS position (*y-axis*) for non PITA (**A**) and PITA (**B**) genes. (**C**) Mean elongation velocities (*y-axis*) across bins of gene length for non PITA and PITA genes. Error bars indicate standard error of the mean. **(D)** Pearson's R values for the pairwise correlations between the relative usage (Ψ) of a gene's AFEs and ALEs based on their ordinal position in mouse expressing wild type RNA pol II (n=110 genes; *left*) and a slow RNA pol II mutant (R749H, n=99 genes, *right*).
